# A polarity-controlled Tad nanomachine enables prey invasion in a bacterial predator

**DOI:** 10.64898/2026.07.08.737194

**Authors:** Coralie Tesseur, Remi Denise, Yoann G. Santin, Géraldine Laloux

## Abstract

Type IV pili are dynamic surface appendages assembled by envelope-spanning nanomachines that mediate diverse bacterial interactions, yet how these systems are adapted to predatory lifestyles remains poorly understood. Here we investigate the tight adherence (Tad) machinery of the obligate predator *Bdellovibrio bacteriovorus.* Using inducible CRISPR interference combined with live-cell and microfluidics imaging, we show that the Tad system is essential for prolonged prey attachment and subsequent prey remodeling and invasion. The machinery assembles specifically at the invasive cell pole before prey encounter and is temporally coordinated with the predatory cell cycle. We further demonstrate that polar localization of the Tad machinery depends on the polarity hub RomR. In addition, the atypical TadZA fusion ATPase interacts with RomR, identifying a potential molecular link between polarity control and Tad assembly. Together, our findings reveal how spatiotemporal control of a conserved filament system supports bacterial predation.

## INTRODUCTION

Bacteria deploy specialized molecular devices to sense and interact with their environment. Type IV pili (T4P) are widespread dynamic filaments of the type IV filament (TFF) superfamily that mediate diverse functions including motility, adhesion, DNA uptake and intercellular interactions. Among T4P, tight-adherence (Tad) pili constitute an evolutionarily distinct subgroup increasingly recognized as versatile nanomachines adapted to diverse bacterial lifestyles ^1–3^. Yet, despite their broad distribution across bacteria, the functional diversification of Tad systems has been explored in only a few model organisms. In *Caulobacter crescentus*, where the Tad system is currently best understood, Tad pili assemble exclusively at the new cell pole in a cell cycle-dependent manner, and cycles of pilus extension and retraction couple surface sensing to developmental transitions ^4–6^. More recently, a Tad-like apparatus named the Kil system was shown to mediate contact-dependent prey killing by the facultative predator *Myxococcus xanthus* ^7–9^, extending the functional repertoire of Tad systems to bacterial predation.

Tad nanomachines differ substantially from canonical T4P in both architecture and regulation (**Figure 1A**). A single bifunctional motor ATPase (TadA) drives extension and retraction of the pilus fiber (a polymer of type IV pilin subunits) through the outer membrane secretin pore, unlike other T4P systems such as the well characterized type IVa pilus (T4aP) that use dedicated motors for these opposite actions ^5,10^. In several bacteria, Tad assembly is spatially restricted to specific subcellular locations that are crucial for function. A ParA/MinD-like ATPase uniquely found in Tad systems (TadZ) promotes unipolar localization of the Tad machinery in *C. crescentus* and is required for TadA polar positioning in *Aggregatibacter actinomycetemcomitans* ^11–15^. In contrast, the Kil system assembles only at prey-contact sites along the *M. xanthus* cell body through unknown spatial determinants ^7–9^. Collectively, these findings indicate that adaptation of Tad systems to distinct developmental programs and behaviors, including predation, relies on precise spatiotemporal control of their assembly and activity.

**Figure 1:**
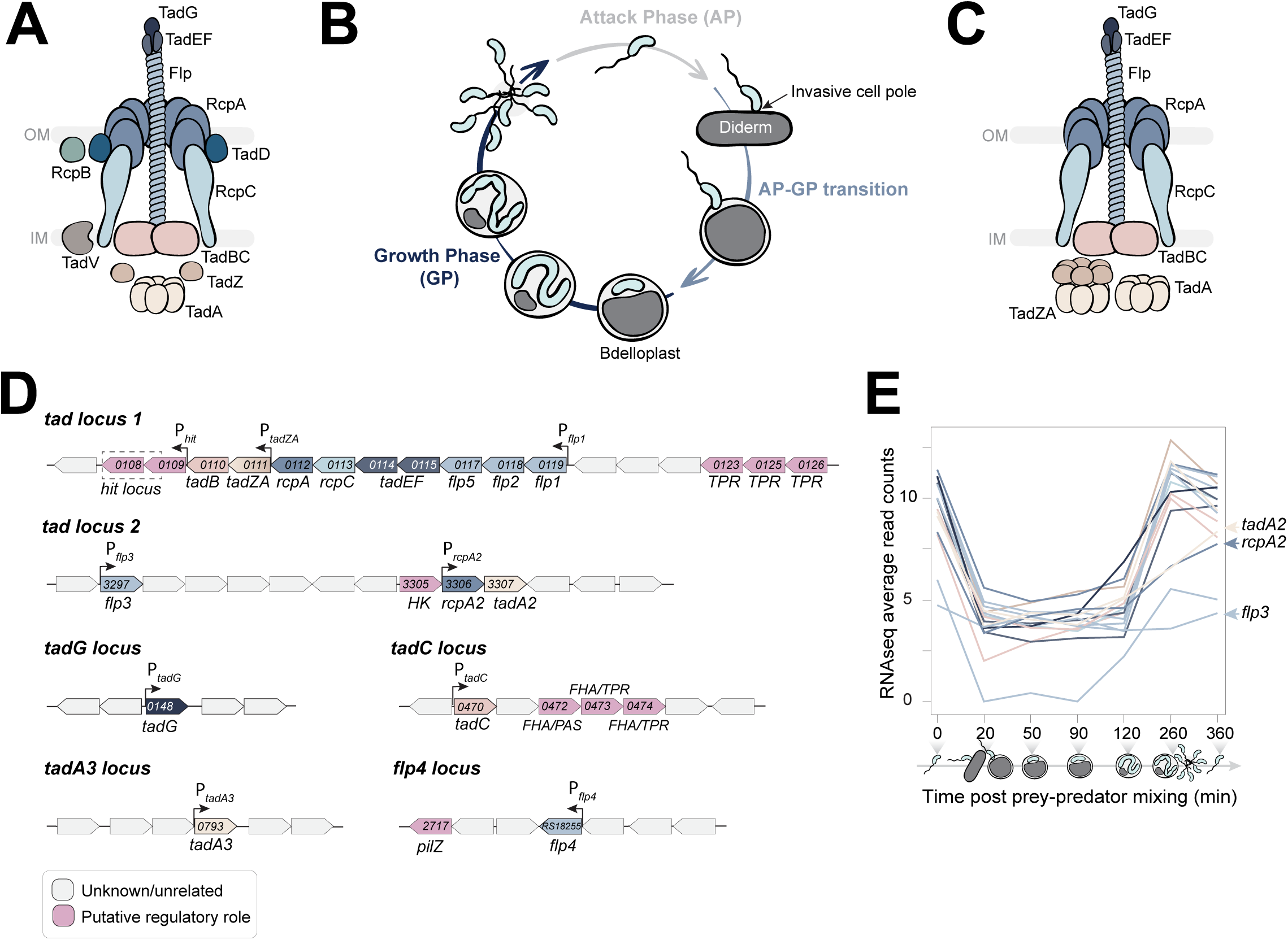
*B. bacteriovorus* encodes a complete Tad machinery. **A.** Molecular architecture of a canonical Tad pilus system according to current models. Tad components are annotated with the Tad/Rcp nomenclature and positioned based on their known or predicted localization within the macromolecular complex. Core components: motor ATPase TadA, prepilin peptidase TadV, IM platform proteins TadB-C, major pilin Flp, minor pilins TadEF, and secretin RcpA. Non-core Tad-specific elements: ParA/MinD-like ATPase TadZ, alignment protein RcpC, pilotin TadD, minor pilin-like protein TadG, and outer membrane-associated protein RcpB. IM = inner membrane; OM = outer membrane. **B.** Schematics of the *B. bacteriovorus* predatory lifecycle. In attack phase (AP), *B. bacteriovorus* swims at high-speed until encountering and attaching to a suitable diderm prey with its invasive cell pole. Then, the predator modifies the prey cell envelope resulting in cell rounding and invades the prey periplasm. The invaded prey is named bdelloplast. There, *B. bacteriovorus* initiates the growth phase (GP) during which it grows as a filament to eventually divide in a variable number of daughter cells, which are released upon prey lysis. **C.** Representation of the predicted Tad machinery of *B. bacteriovorus*. The color code and annotations are kept consistent with **A** for comparison. IM = inner membrane and OM = outer membrane. **D.** Annotations and genetic organization of the six *tad* loci identified in the *B. bacteriovorus* HD100 genome. *tad* genes are colored according to their predicted protein product within the complex depicted in **C**. Predicted promoters (P*_x_*) are indicated with arrows. Gene identifiers are indicated without the ‘Bd’ prefix. *hit locus*: “host interaction” locus proposed to play a role in T4P regulation. The represented boundaries were selected arbitrarily to illustrate the genetic environment of *tad* genes and the presence of neighboring genes coding for proteins with putative regulatory functions, for which selected protein domains are indicated. TPR = tetratricopeptide repeats; FHA = forkhead-associated; PAS = Per-ARNT-Sim; HK = histidine kinase. **E.** *tad* genes are specifically expressed during early predation stages. Expression profiles of *tad* genes were recovered from a time-resolved RNA-sequencing (RNA-seq) dataset ^1^, in which mRNA levels were assessed at 0, 20, 50, 90, 120, 260, 360 min post prey-predator mixing. For each timepoint, the corresponding cell cycle stage is depicted. Curves correspond to the different *tad* genes, colored according to **D**. Arrowheads highlight genes that display distinct expression profiles (i.e., *tadA2, rcpA2, flp3* and *flp4*).

*Bdellovibrio bacteriovorus* is an obligate predatory bacterium that proliferates within the periplasm of other diderm bacteria. During its lifecycle, *B. bacteriovorus* alternates between a swimming, non-replicative attack phase and an intra-periplasmic growth phase within the killed and modified prey, termed bdelloplast (**Figure 1B**) ^16^. Filamentous growth and multiple asynchronous rounds of DNA replication, followed by non-binary cell division, generate a variable number of progeny that exit the prey to resume the attack phase ^17–19^. How predators recognize, attach to and invade their prey is poorly understood, although these early predation stages are essential for cell cycle progression ^16,20^. Previous studies suggest that the invasive cell pole – the pole by which predators interact with prey – constitutes a highly specialized platform enriched in signaling networks and macromolecular complexes, with the polarity protein RomR proposed to act as a central organizer ^21–23^. Consistent with this idea, cryo-electron tomography revealed multiple trans-envelope assemblies decorating the invasive pole, including structures reminiscent of T4P machineries and extracellular filaments ^24^. Yet, the identity and function of these complexes remain unclear.

The *B. bacteriovorus* genome is predicted to encode both a T4aP and a Tad system, although the composition and organization of the Tad machinery remain incompletely resolved ^2,9,25–27^. Expression of Tad pilin genes peaks during the attack phase, and several appear essential for predation ^26,28^. Intriguingly, the major *tad* cluster lies adjacent to the so-called “host-interaction” (*hit*) locus, whose mutation can promote some prey-independent proliferation and alter piliation ^29–31^. Although the molecular function of the *hit* locus and its precise link with pili systems remains undetermined, these observations suggest that the Tad machinery may be connected to pathways regulating predatory behavior ^28,31^. In addition, the predicted *B. bacteriovorus* machinery encodes an atypical TadA-TadZ fusion protein ^13^, hinting at a potential coupling between Tad activity and spatial regulation. However, the Tad system has never been studied functionally in wild-type predatory *B. bacteriovorus*, largely because the obligate nature of predation prevents genetic inactivation of systems required for predation.

Here we combined CRISPR interference with live fluorescence and microfluidics microscopy to directly investigate the role and spatiotemporal dynamics of the Tad system during the predatory lifecycle of *B. bacteriovorus*. We show that an intact Tad machinery is essential for prolonged attachment, prey remodeling and invasion, and that it localizes at the invasive cell pole during these key predation stages. Furthermore, we demonstrate that Tad positioning depends on the invasive pole organizer RomR, which interacts with the TadZ domain of the atypical Tad ATPase. Together, our findings reveal how polarity cues promote assembly of an essential Tad nanomachine during bacterial predation.

## RESULTS

### *Bdellovibrio bacteriovorus* carries a complex genetic repertoire encoding a complete Tad machinery

To establish a comprehensive inventory of Tad components in the reference strain *Bdellovibrio bacteriovorus* HD100, we performed a genome-wide analysis using a MacSyFinder-based pipeline ^1,32^ refined by targeted homology searches and domain architecture analysis (Methods). This analysis confirms that the *B. bacteriovorus* genome encodes a complete Tad nanomachine. All conserved core components characteristic of the TFF superfamily are present ^1^, including major (Flp) and minor pilins (TadEF), an outer membrane secretin (RcpA), inner membrane (IM) platform proteins (TadB and TadC), and a cytoplasmic ATPase (TadA), together with Tad-specific accessory factors such as RcpC, TadG and TadZ (**Figure 1A, C-D**).

However, the *B. bacteriovorus* Tad repertoire differs from canonical systems in several ways. First, the genomic organization of *tad* genes is unusually dispersed. While most core genes cluster within a primary locus (hereafter *tad locus 1*), TadC is encoded separately (*tadC locus*), unlike in other species. Moreover, additional *tad* genes are distributed across at least four other genomic regions, including a duplicated *rcpA-tadA* module (*tad locus 2*), a third isolated *tadA* paralog (*tadA3* locus), and a lone *tadG* encoding a pilus tip protein ^9^. Strikingly, *tad locus 1* encodes an atypical TadZA hybrid protein in which the ParA/MinD-like protein TadZ is fused to the canonical TadA ATPase.

Another unusual feature of the *B. bacteriovorus* Tad system is the apparent absence of canonical TadV prepilin peptidase within *tad* loci ^26^. Because pilin subunits must be processed from prepilins before incorporation into polymerizing filaments, we tested whether prepilin processing could instead be shared between distinct TFF systems, as reported in a limited number of bacteria ^1^. Consistent with this idea, we found that the T4aP-associated PilD prepilin peptidase, encoded elsewhere on the chromosome, efficiently processed both the major T4aP pilin PilA and the major Tad pilin Flp1 when heterologously produced in *E. coli*, whereas PilD catalytic mutants did not (**Figure S1**).

Beyond structural components, several *tad* genes are located near genes encoding predicted regulatory proteins harboring tetratricopeptide repeats (TPR) domains (mediating protein-protein interactions), forkhead-associated (FHA) domains (involved in phosphorylation-dependent signaling), PilZ domains (c-di-GMP binding) and histidine kinase domains (**Figure 1D**). Collectively, these analyses reveal that *B. bacteriovorus* encodes a structurally complete yet genetically dispersed Tad nanomachine, which is potentially integrated in broader signaling networks.

### Genes encoding the core Tad machinery are essential and expressed during early predation stages

We first asked whether the Tad machinery is required for the predatory lifecycle of *B. bacteriovorus*. Our attempts to delete core *tad* genes (*rcpA, tadZA, tadB* or *tadC*) were unsuccessful (see Methods), and genes encoding predicted pilins *flp1, flp2* and *flp4* could not be deleted in a previous study ^26^. This shows that these components are essential for survival. In contrast, deletion of the duplicated *rcpA2-tadA2* module did not impair predation, showing that these paralogs are dispensable for Tad function under the tested conditions (**Figure S2**). These observations indicate that the core Tad machinery is essential in *B. bacteriovorus*.

We next examined the expression of *tad* genes. Previous studies reported attack phase-specific transcription of *flp* pilin genes ^26,28^, and both the RcpA secretin and pilin Flp1 were detected in attack-phase predator proteomes ^33,34^. Consistent with these observations, our time-resolved RNA-seq analysis across a synchronized predatory cycle ^35^ revealed that expression of most *tad* genes peaks when progeny is released from prey and during the subsequent attack phase. By contrast, *flp3* and the duplicated *rcpA2-tadA2* module displayed distinct expression profiles, suggesting accessory or Tad-unrelated functions (**Figure 1E**). Hence, essential *tad* genes are maximally expressed when predators encounter and invade prey, supporting a key role for the Tad machinery during early predation.

### The Tad machinery is required for predation

Because the Tad system could not be genetically inactivated, we used inducible CRISPR interference (CRISPRi) to conditionally deplete selected Tad components in *B. bacteriovorus*. Here, chromosomal *dcas9_Spa_* expression is induced by IPTG ^36^, whereas a constitutively expressed plasmid-encoded sgRNA targets either (i) the *tadZA-tadB* operon, encoding the Tad ATPase fusion protein and an inner membrane platform protein, (ii) the secretin gene *rcpA*, or (iii) the minor pilin gene *tadG*. CRISPRi induction during overnight predatory cultures caused severe predation defects for all three targets: cultures remained turbid, indicating failure to clear the *E. coli* population, unlike non-induced conditions or a non-targeting sgRNA control **(Figure 2A)**. Phase contrast microscopy confirmed the low predator numbers together with abundance of intact *E. coli* cells upon *tad* depletion **(Figure 2A)**. These results demonstrate that the Tad machinery is required for efficient predation at the population level. We next investigated how Tad depletion affects early prey-predator interactions. To selectively analyze the predatory capacity of newborn Tad-depleted *B. bacteriovorus*, CRISPRi was induced during a single synchronized round of intra-prey growth, as described previously ^36^. Newly released attack-phase progeny were then mixed with prey, and bdelloplast formation was monitored over time using *E. coli* cell rounding as a proxy. Silencing *tadZA-tadB*, *rcpA* or *tadG* strongly reduced the proportion of *E. coli* cells converted into bdelloplasts relative to non-targeting or uninduced controls (**Figure 2B**), showing that an intact Tad machinery is required for prey invasion.

**Figure 2:**
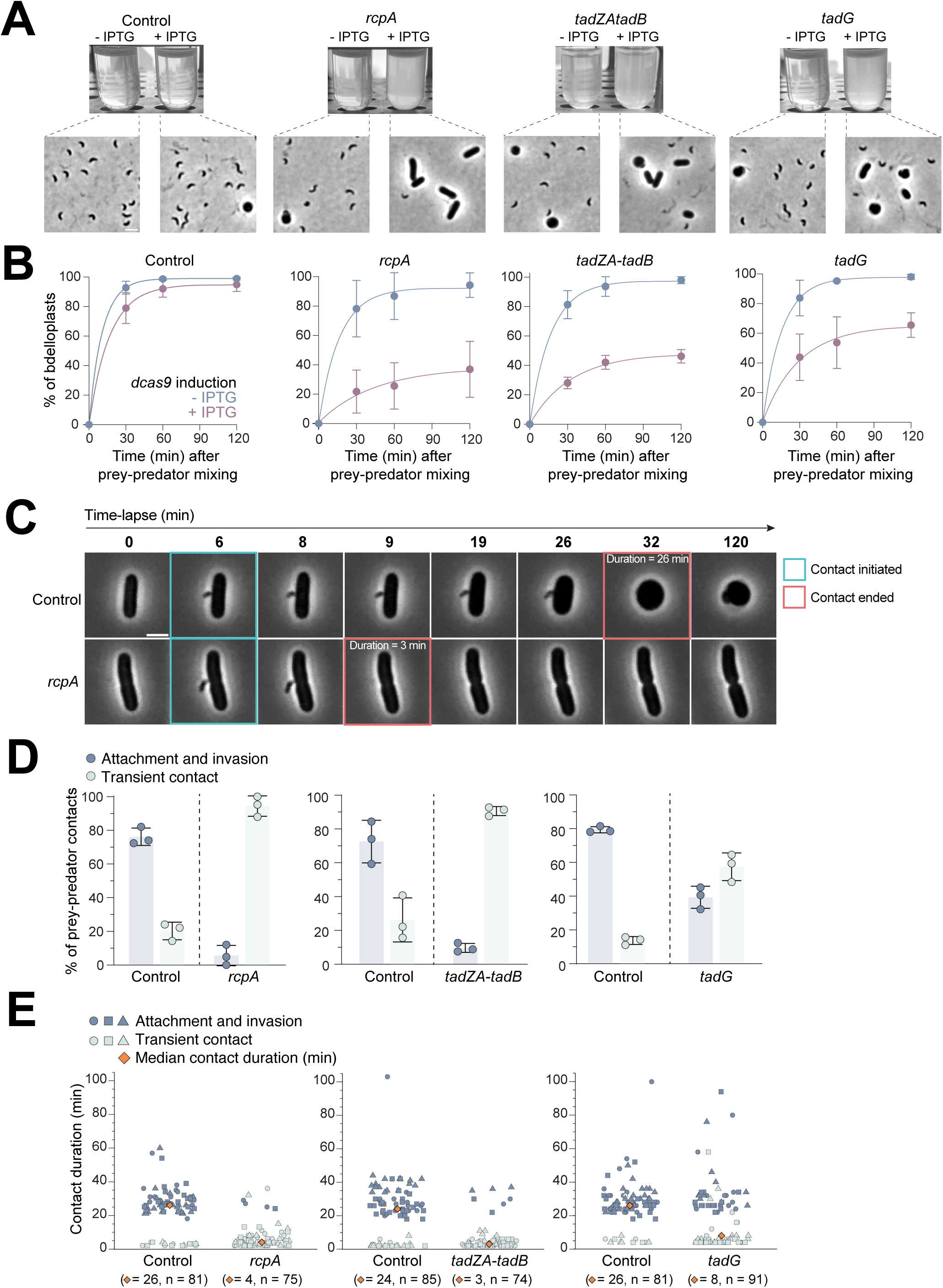
An intact Tad machinery is required for prey attack. Tad-associated phenotypes were assessed with CRISPRi strains expressing sgRNAs for the targeted repression of *rcpA* (GL2000 *rcpA*sgRNA), *tadZA-tadB* (GL2000 *tadZA-tadB*sgRNA), and *tadG* (GL2914). For clarity, only the targeted gene is indicated in each panel. The control condition corresponds to a strain expressing a non-targeting sgRNA (GL2915). **A.** Overnight repression of *rcpA*, *tadZA-tadB* or *tadG* (GL2914) results in reduced predator proliferation. Top panel: pictures of overnight prey-predator co-cultures incubated in absence or presence of 200 µM IPTG as indicated for induction of *dcas9* expression. Bottom panel: representative phase contrast images of the corresponding overnight cultures. Scale bar = 2 µm. **B.** Quantification of bdelloplast formation over time after mixing newborn *B. bacteriovorus* – obtained after a single round of synchronized growth in absence or presence of 200 µM IPTG for *dcas9* induction – with exponentially grown *E. coli* cells. Phase contrast microscopy snapshots were acquired in a time-course experiment at 0, 30, 60 and 120 min post prey-predator mixing, and *E. coli* roundness was used as a proxy for bdelloplast formation. Dots and errors bars represent the mean and standard deviations of at least three independent biological replicates. Data and number of cells analyzed are available in **Table S1,** which will be included in the peer-reviewed version of the manuscript. **C-E.** After a single synchronized predation cycle in presence of 200 µM IPTG to induce *dcas9* expression, newborn Tad-depleted and control predators were perfused in separate microfluidics chambers containing immobilized exponentially grown *E. coli*. Early stages of predation were monitored by time-lapse phase contrast microscopy with 1-min (RcpA and TadZA-TadB depletion) or 2-min (TadG depletion) intervals for 2h. **C.** Representative phase contrast images of prey-predator interactions at selected timepoints of time-lapses in control (top) or RcpA-depleted conditions (bottom). Time points at which contact initiates (blue squares) and ends (red squares), as well as the resulting contact duration, are indicated for each condition. **D.** Quantification of the fraction of contacts (visible for at least 2 consecutive time points) either resulting in prey attachment and invasion or followed by detachment from prey (transient contact). Bars, error bars, and dots represent the mean, standard deviations and replicates values of three independent biological replicates, respectively. Detailed percentages and numbers of cells analyzed are available in **Table S1,** which will be included in the peer-reviewed version of the manuscript. **E.** Scatter plot of contact durations measured in **D.**, distinguishing events leading to prey attachment from transient contacts (i.e., detachment from the prey cell surface). Circles, squares, and triangles denote three independent biological replicates. Medians and number of contacts analyzed (n) are indicated.

### Tad depletion causes transient prey attachment without remodeling or invasion

Time-lapse imaging in microfluidic chambers revealed that Tad-depleted *B. bacteriovorus* fail to achieve early stages of prey interaction. In this system, attack-phase predators were allowed to freely swim, encounter and invade immobilized prey. We monitored prey-predator contacts, considering only those that persisted for more than one time point to exclude random collisions, as well as prey invasion, defined as full enclosure of the *B. bacteriovorus* cell within the prey. Whereas most contacts led to prey invasion under control conditions, they rarely progressed to invasion by Tad-depleted predators. Instead, the *B. bacteriovorus* cells detached and resumed swimming (**Figures 2C-D and S3A**). Moreover, these transient contacts were shorter than the attachment phase preceding prey invasion by control predator cells (**Figure 2E**). Remarkably, *E. coli* cells contacted by Tad-depleted *B. bacteriovorus* never underwent the characteristic rounding associated with bdelloplast formation. Furthermore, these cells continued elongating and occasionally divided during the experiment (**Figures 2C and S3B-C**). Predation defects were consistently stronger upon depletion of core components (TadZA-TadB and RcpA) than upon depletion of the minor pilin TadG (**Figure 2A-D**), either due to a less efficient dCas9 targeting by the sgRNA ^37,38^ or reflecting the enhancing yet accessory role of TadG in filament polymerization reported in other species ^39,40^. Altogether, our results show that a complete Tad machinery is necessary for prolonged attachment to prey and subsequent prey remodeling and invasion.

### The Tad machinery localizes at the invasive cell pole in a cell cycle-dependent manner

Because early predation functions involve the invasive pole, we investigated whether the Tad machinery assembles at this specialized subcellular region. We used the secretin RcpA as a main reporter for Tad assembly, as established in other Tad systems ^8,11^. To preserve the native *tad locus 1* (**Figure 1D**) and maintain a pool of untagged secretin subunits, we introduced a second chromosomal copy encoding an RcpA-mCherry fusion under control of its native promoter at an ectopic locus in *B. bacteriovorus*. Immunoblot analysis confirmed production of the full-length fusion protein (**Figure S4A**), whose abundance fluctuated during the cell cycle in agreement with the native *rcpA* transcriptional profile (**Figure 1E**, **Figure S4B**). Moreover, the resulting merodiploid strain displayed normal morphology and predatory efficiency relative to wild-type cells (**Figure S4C-D**), indicating that the fluorescent reporter does not impair cell physiology or Tad function.

In attack-phase cells, RcpA-mCherry formed a single polar focus even in the absence of prey, and demograph analysis confirmed strong signal enrichment at one cell pole (**Figure S4E**). The inner membrane platform protein TadC, tagged with mCherry and produced from a plasmid under control of the native *P_tadC_* promoter, also formed unipolar foci. Imaging RcpA-mCherry or TadC-mCherry together with the invasive-pole marker RomR-msfGFP ^23^ revealed robust co-localization at this pole for both Tad components (**Figure 3A, Figure S4C-D**,**F-G).** Early prey-predator contacts imaged shortly after mixing further showed that RcpA-mCherry remains associated with the invasive pole during prey attachment (**Figure S4H**). Altogether, these observations indicate that the Tad machinery is assembled at the invasive pole before prey encounter and remains positioned there during contact with prey, consistent with its essential role in early predation.

**Figure 3:**
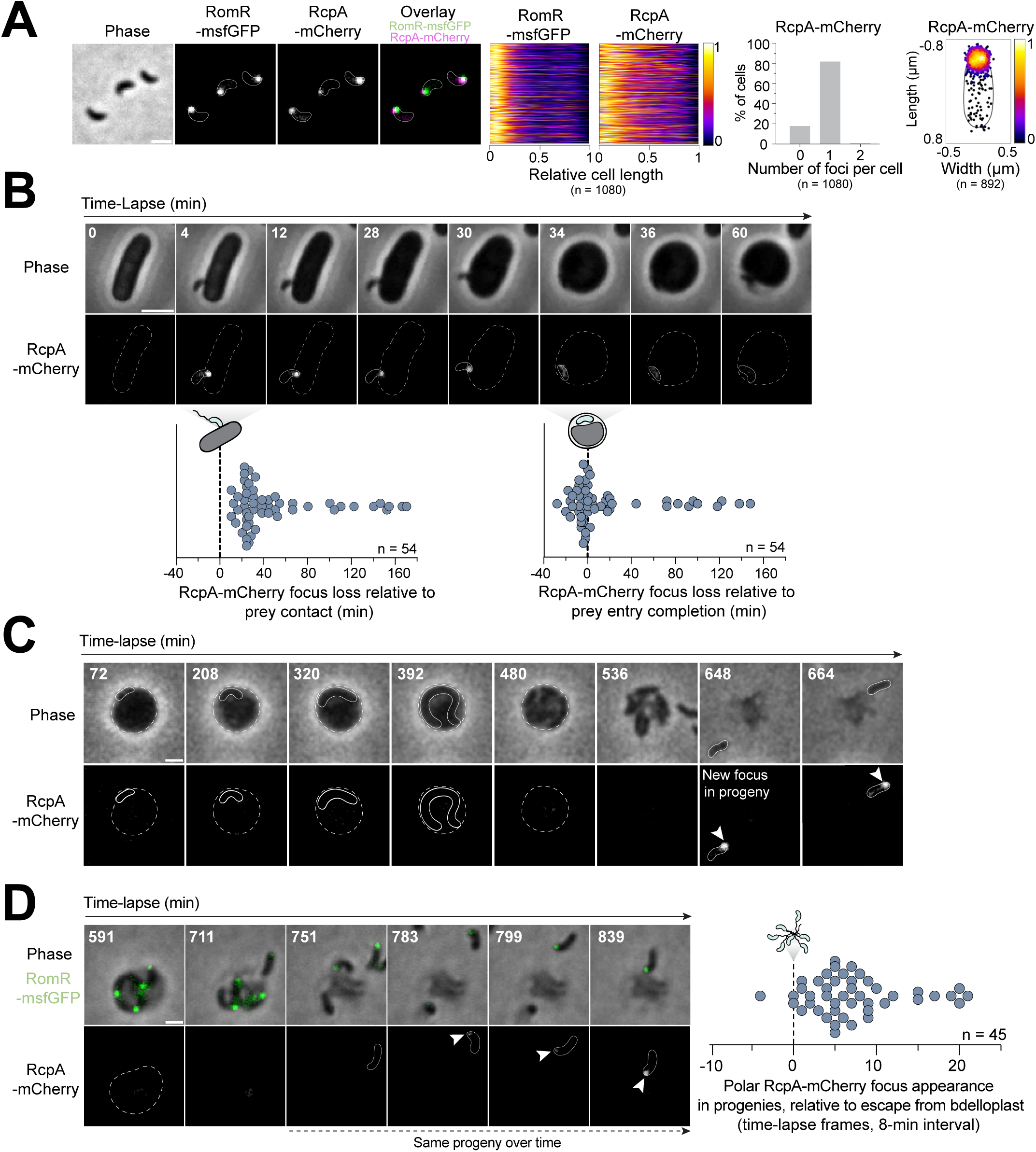
The Tad machinery localizes at the invasive pole in a cell cycle-dependent manner. **A.** Left to right: representative phase contrast and epifluorescence images of attack phase *B. bacteriovorus* cells producing RcpA-mCherry and the invasive cell pole marker RomR-msfGFP (*romR::romR-msfgfp Bd0063-Bd0064::*P*_flp1_-rcpA-mcherry*; strain GL2585); demographs showing signals distribution along the relative cell length for cells oriented based on the RomR-msfGFP signal intensity; histogram of the number of RcpA-mCherry foci per cell; 2D heatmap of the subcellular localization of RcpA-mCherry foci in cells normalized by the cell length and oriented based on RomR-msfGFP signal intensity. n corresponds to the number of cells analyzed in a representative experiment from three independent biological replicates. **B.** Time-lapse microscopy imaging (2-min interval) of prey-predator interactions upon perfusing *B. bacteriovorus* producing RcpA-mCherry (GL2479) in a microfluidic chamber in which exponentially grown *E. coli* cells are immobilized. Top: representative phase contrast and epifluorescence images of selected timepoints in a representative experiment. Bottom: monitoring of RcpA-mCherry focus loss relative to prey contact (left) and prey entry completion (right). n indicates the number of contacts analyzed from two independent biological replicates. All analyzed predators were seen to progress to the growth phase (i.e. elongation within the bdelloplast). **C-D.** RcpA-mCherry is absent during the growth phase and reappears after escape from the bdelloplast. **C.** Representative phase contrast and epifluorescence images of a bdelloplast at selected timepoints of a time-lapse experiment. Attack phase *B. bacteriovorus* cells producing RcpA-mCherry (GL2479) were mixed with prey for 72 min prior to imaging at 8-min intervals. This time-lapse experiment was performed twice. **D.** Left: representative phase contrast and HiLo fluorescence images of a bdelloplast in late growth phase at selected timepoints of a time-lapse experiment. Attack phase *B. bacteriovorus* cells producing RcpA-mCherry and RomR-msfGFP (GL2585) were mixed with prey for 71 min prior to imaging at 8-min intervals. Right: monitoring of the timing of reappearance of polar RcpA-mCherry foci colocalizing with RomR-msfGFP in progenies, relative to escape from bdelloplast. n denotes the number of cells analyzed from two independent biological replicates. Scale bars = 1 µm (**A**, **C** and **D**) or 2 µm (**B**).

Time-lapse microscopy allowed us to monitor Tad dynamics throughout the predatory cycle (**Figure 3B-D**). RcpA-mCherry remained stably localized at the invasive pole during the initial attachment phase but disappeared shortly after prey invasion started (**Figure 3B**). During filamentous intra-prey growth, RcpA foci were usually absent (98.81% cells, n = 84, **Figure 3C**) and reappeared only after cell division and progeny release, as freshly escaped attack-phase cells again displayed a single polar RcpA-mCherry focus (**Figure 3C-D**). Co-imaging with RomR-msfGFP confirmed that this focus associates with the invasive pole (**Figure 3D**). The disappearance of RcpA upon prey invasion and its reappearance in progeny cells are consistent with the observed cell cycle-dependent fluctuations in protein levels (**Figure S4B**). Altogether, our data demonstrate that Tad localization is tightly coordinated with the predatory cell cycle: the machinery is assembled at the invasive pole in attack-phase cells, lost upon prey entry, absent during intra-prey growth, and quickly re-established in the progeny before the next round of predation.

### RomR is required for invasive pole localization of the Tad machinery

RomR is an essential cytosolic protein that marks future invasive poles before cell division and was previously linked to a network involving T4P-associated proteins, including TadZA ^23^. We therefore tested whether RomR controls polar positioning of the Tad machinery. Since perturbation of chromosome segregation by *parB* overexpression results in cell division and polarity defects ^17,41^, we examined RcpA and RomR localization in these conditions. Elongated *parB*-overexpressing cells indeed displayed aberrant RomR positioning, including bipolar or non-polar foci (**Figure 4A**). Strikingly, RcpA-mCherry mirrored RomR localization patterns and we rarely observed RcpA polar localization independent of RomR (**Figure 4A**). These observations are consistent with the idea that positioning of the Tad primarily follows RomR-defined cues.

**Figure 4:**
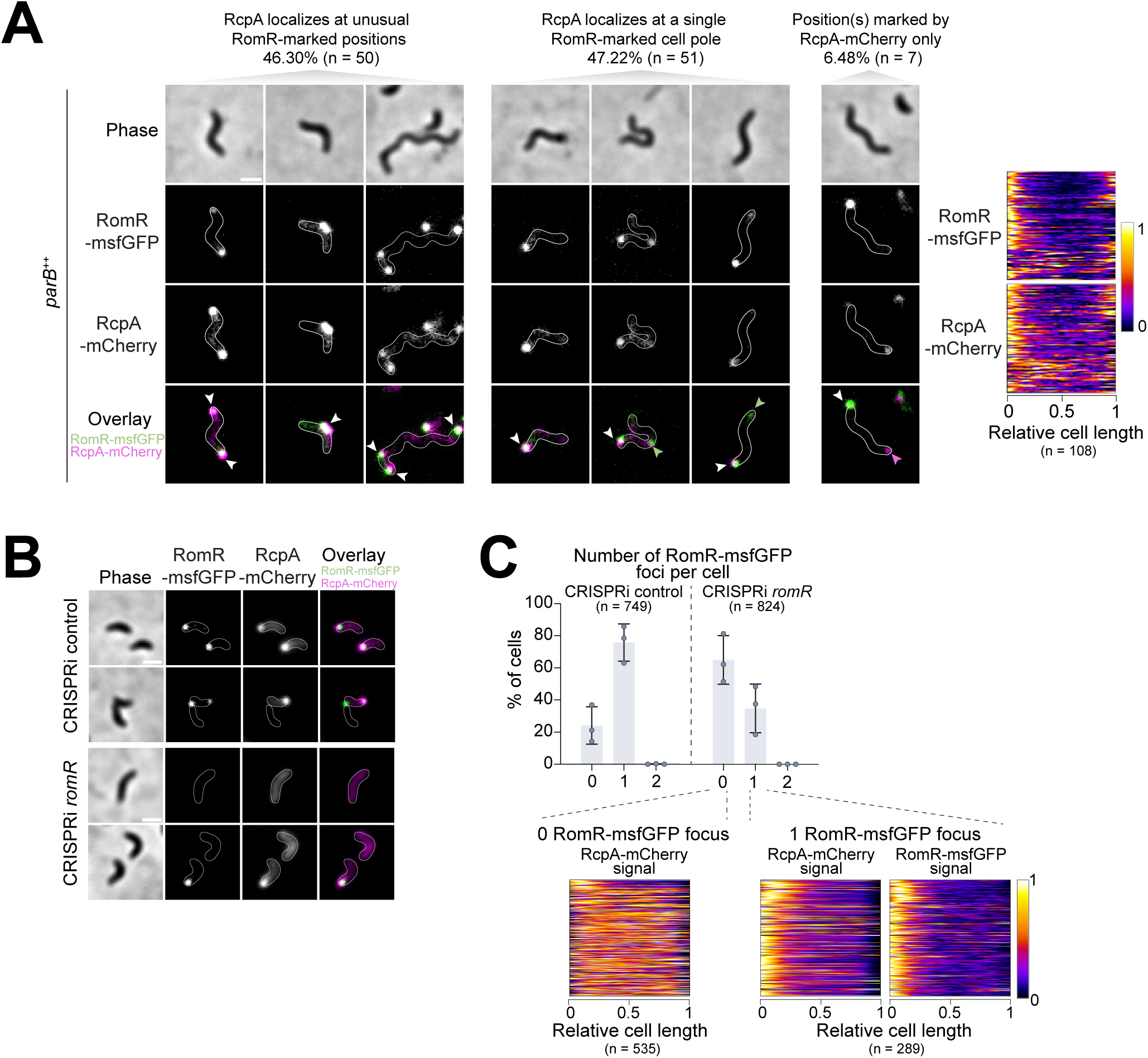
RomR contributes to the proper localization of the RcpA secretin. **A.** Co-localization of RomR-msfGFP and RcpA-mCherry in cells constitutively overexpressing *parB* (*parB^++^*) from a replicative plasmid (strain GL2585 *parB^++^*) and displaying abnormal cell lengths (≥ 1.7 µm). Left: representative phase contrast and epifluorescence images of attack phase cells. Cells were grouped according to their RcpA/RomR co-localization patterns as indicated. Green, magenta, and white arrowheads indicate RomR-msfGFP, RcpA-mCherry, and co-localizing RomR-msfGFP/RcpA-mCherry foci, respectively. Right: demographs showing the distribution of RomR-msfGFP and RcpA-mCherry signals. n indicates the number of cells analyzed from 3 independent biological replicates. **B-C.** Localization of RcpA-mCherry upon CRISPRi-mediated depletion of RomR-msfGFP. CRISPRi strains expressing a non-targeting control sgRNA (CRISPRi control, strain GL3343 control sgRNA**, B**) or a sgRNA targeting *romR* (CRISPRi *romR*, strain GL3343 *romR*sgRNA, **C**) were grown for a single synchronized cycle in presence of 200 µM IPTG to induce *dcas9* expression. **B.** representative phase contrast and epifluorescence images of newborn attack phase cells obtained upon IPTG addition during one growth cycle in control (top) or RomR-depleted conditions (bottom). **C.** Top: RomR depletion results in a decrease in the number of RomR-msfGFP foci per cells. Histograms of the number of RomR-msfGFP foci per cells. The number of cells (n =) analyzed from three independent biological replicates are indicated. Bottom: RcpA-mCherry polar localization is disrupted in absence of a detectable RomR-msfGFP focus. Demographs show the distribution of RcpA-mCherry signal (left) or RcpA-mCherry and RomR-msfGFP signals (right) in cells displaying 0 or 1 RomR-msfGFP focus, respectively. Cells were oriented based on the RomR-msfGFP signal intensity distribution. n corresponds to the number of cells analyzed from three independent biological replicates. **A-B** scale bars = 1 µm.

We next directly tested whether RomR is required for Tad localization using CRISPRi depletion in cells carrying both the RcpA-mCherry reporter and a *romR-msfgfp* fusion replacing the native *romR* gene (**Figure S5A**). RomR depletion during a single growth cycle or overnight predatory culture caused severe predation defects as indicated by reduced bdelloplast formation (**Figure S5B**). Time-lapse imaging in microfluidics showed that RomR is required for initial prey attachment. When an equal mix of RomR-depleted and control populations were perfused in the chamber (**Figure S5C**), RomR-depleted cells contributed to only 18±10% of contacts (**Figure S5D**). Moreover, RomR-depleted cells displayed slightly more variable cell length compared to wild-type (**Figure S5B,E**), likely reflecting pleiotropic effects due to the removal of an essential hub protein.

Strikingly, in most cells lacking a detectable RomR-msfGFP signal, RcpA-mCherry no longer formed polar foci and instead appeared diffuse **(Figure 4B-C, Figure S5F)**. Conversely, cells retaining RomR signal – due to incomplete depletion – maintained polar RcpA-mCherry foci co-localizing with RomR-msfGFP (**Figure 4B-C, Figure S5F**). Together, these results demonstrate that RomR is required for localization of the Tad machinery at the invasive pole in *B. bacteriovorus*.

### The TadZA fusion ATPase interacts with RomR and localizes dynamically in a RomR-dependent manner

The presence of a TadZA fusion protein **(Figure 1C and 5A)** suggests potential coupling between TadA-dependent pilus dynamics and spatial regulation of the Tad nanomachine. TadZA comprises an N-terminal ParA/MinD-like TadZ region containing a Walker A-like motif, connected by a flexible linker to the C-terminal TadA AAA ATPase (**Figure 5A**). Since TadZA was previously identified within the RomR proximity network ^23^, we investigated whether both proteins interact directly. POLAR assays in *E. coli* ^42,43^ revealed robust recruitment of TadZA-mScarlet by pole-tethered RomR-msfGFP-H3H4 (**Figure 5B**). Using truncated TadZA variants revealed that the TadZ region together with the linker is necessary and sufficient for interaction with RomR, whereas neither TadZ nor TadA alone, nor TadA combined with the linker, supported recruitment (**Figure 5B**). Because *E. coli* lacks homologs of the RomR polarity network and Tad components, polar recruitment in this heterologous context likely reflects a direct interaction between RomR and the TadZ portion of TadZA.

**Figure 5:**
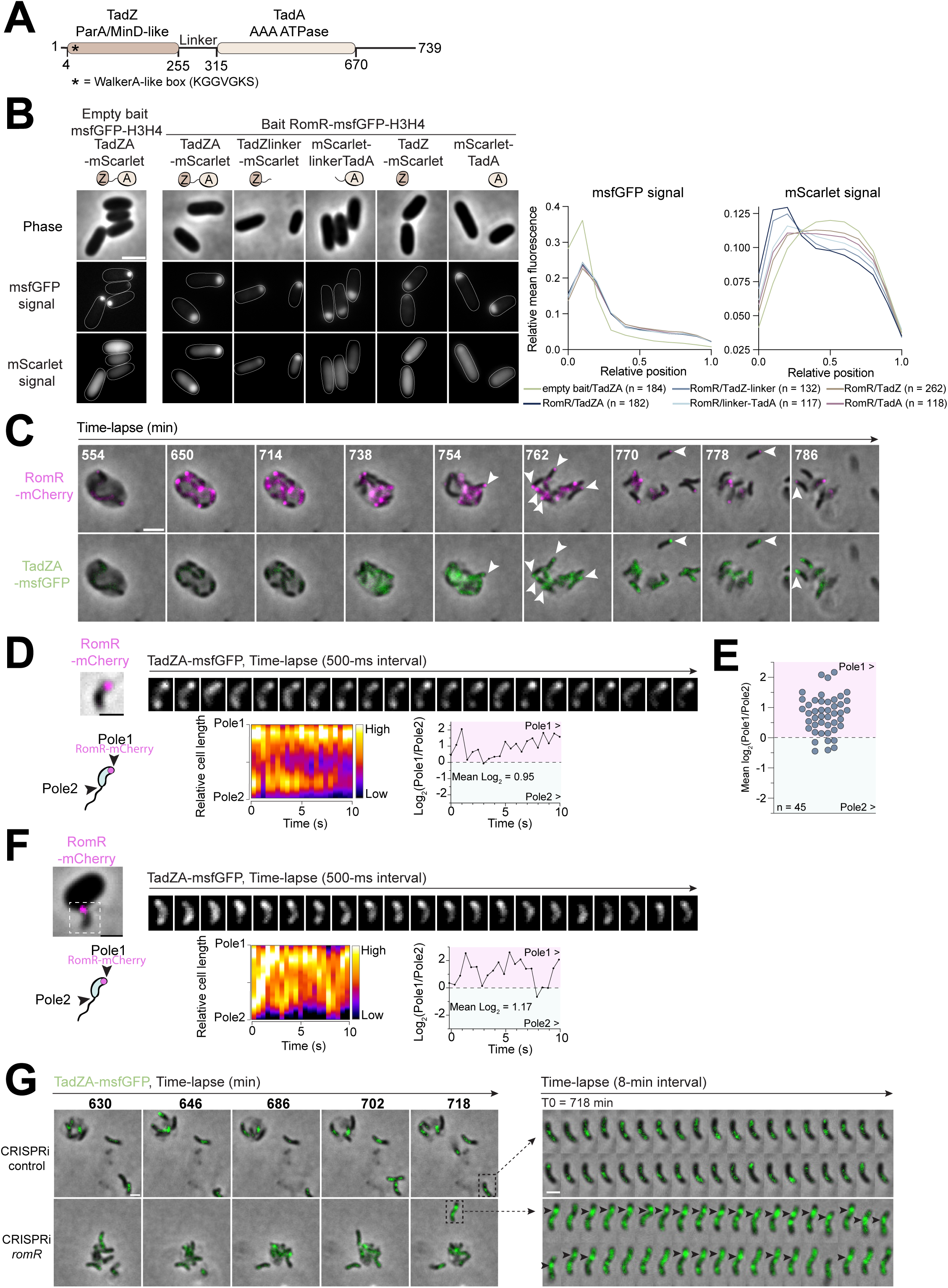
TadZA interacts with RomR and displays a dynamic localization pattern in attack phase. **A.** Schematic representation of the domain architecture of the *B. bacteriovorus* TadZA fusion ATPase. The N-terminal TadZ region contains the WalkerA-like motif (consensus K/RGGXGXS/T). **B.** RomR and TadZA interact in POLAR assays. Left: representative phase contrast and epifluorescence images of the tested combinations of bait (RomR-msfGFP-H3H4 or msfGFP-H3H4 alone) and test (mScarlet-tagged) protein fusions. Right: relative mean fluorescence intensity profiles of msfGFP and mScarlet signals along relative cell length for all POLAR combinations tested. The number of cells analyzed (n) in representative experiments are indicated. **C.** TadZA-msfGFP signal accumulates at invasive cell poles upon bdelloplast escape. Representative phase contrast and HiLo fluorescence images of a bdelloplast in late growth phase, at selected timepoints of a time-lapse experiment. *B. bacteriovorus* attack phase cells producing TadZA-msfGFP and RomR-mCherry (GL3452) were mixed with prey for 74 min prior to imaging at 8-min intervals. White arrowheads indicate polar accumulation of TadZA-msfGFP at the invasive cell pole of progenies marked by RomR-mCherry. **D-E.** TadZA-msfGFP is highly dynamic in attack phase cells, but is enriched at the invasive cell pole. **D.** Top: representative phase contrast and HiLo fluorescence images of a representative *B. bacteriovorus* cell producing TadZA-msfGFP and RomR-mCherry (GL3452). A phase contrast and RomR-mCherry snapshot was acquired before 500-ms interval time-lapse imaging of TadZA-msfGFP. Bottom: schematic representation of the system used for dynamics characterization, where Pole1 and Pole2 correspond to cell areas from each cell extremity to 20% of the cell length. Pole1 is identified as the invasive cell pole by the RomR-mCherry focus. Kymograph representing the distribution of TadZA-msfGFP signal between Pole1 and Pole2 over time in the same cell. Variation of the log_2_(Pole1/Pole2) fluorescence ratio over time, where values > 0 correspond to higher fluorescence intensity at Pole1, values = 0 to equal fluorescence levels at both poles, and values < 0 to higher fluorescence levels at Pole2. The mean log_2_ ratio across the entire time-lapse is indicated. **E.** Dot plot showing the mean log_2_(Pole1/Pole2) fluorescence ratio measured in individual attack phase cells imaged by time-lapse microscopy as in **D.** n indicates the number of cells analyzed from three independent biological replicates. **F.** The TadZA-msfGFP signal remains dynamic during prey attachment. Top: representative phase contrast and HiLo fluorescence images of a representative *B. bacteriovorus* cell producing TadZA-msfGFP and RomR-mCherry (GL3452) attached to a prey cell 20 min after mixing attack phase predators with stationary phase-grown *E. coli*. Phase contrast and RomR-msfGFP snapshots and TadZA-msfGFP time-lapse images were acquired as in **D**. Bottom: Kymograph and quantification of log_2_(Pole1/Pole2) fluorescence ratio overtime as in **D**. **G.** TadZA-msfGFP subcellular localization and dynamics are disturbed in RomR-depleted progenies. *B. bacteriovorus* cells producing TadZA-msfGFP (*pepN-folE::*P*flp1-tadZA-msfgfp*) and expressing a non-targeting control sgRNA (CRISPRi control, top; GL3570) or a sgRNA targeting *romR* (CRISPRi *romR*, bottom; GL3571) were mixed with prey for 130 min prior time-lapse imaging, in presence of 200 µM IPTG to induce *dcas9* expression. To focus on newly generated progenies escaping from bdelloplasts, images were acquired every hour for 7 hours, then every 8-min. Both strains were imaged on two separate areas of the same agarose pad containing 200 µM IPTG to maintain *dcas9* expression during the whole time-lapse. Left: Phase contrast and fluorescence overlays for representative escaping progenies are shown for the indicated time points. Right: 8-min interval images of the cell shown in the corresponding inset to illustrate distinct TadZA-msfGFP subcellular localization and dynamics, with apparent pole-to-pole focus oscillations in the control vs larger, less dynamic, subpolar (black arrowheads) accumulations in RomR-depleted cells. In **B.** and **G**., for display purposes only, brightness and contrast settings of epifluorescence images were adjusted individually, to illustrate protein localization patterns rather than differences in fluorescence intensities. Scale bars = 2 µm (**B** and **C**) or 1 µm (**D**, **F** and **G**). All experiments were performed at least twice.

The RomR/TadZA interaction prompted us to examine TadZA dynamics throughout the predatory cycle. We therefore introduced a second chromosomal copy encoding a functional TadZA-msfGFP fusion under control of the native promoter (**Figure S6A-B**). Accumulation of the full-length TadZA-msfGFP fusion over time was consistent with the RNA-seq expression profile (**Figure 1E and S6C**), and fluorescence was correspondingly low or undetectable during intra-prey growth (**Figure 5C**). Around the time of cell division, TadZA-msfGFP started to accumulate in (future) daughter cells and formed weak foci that mostly colocalized with RomR-mCherry upon escape from the bdelloplast (**Figure 5C**). Then, after prey exit TadZA-msfGFP displayed highly dynamic behavior, unlike the stable invasive-pole localization observed for RcpA, forming foci or larger cloud-like structures that rapidly shifted between both cell poles within less than a second (**Figure 5D**). Yet, quantitative analysis of fluorescence between both poles revealed an enrichment of the TadZA-msfGFP signal at the RomR-labelled invasive pole over time (**Figure 5D-E**). This asymmetric distribution remained unchanged upon prey contact (**Figure 5F**) and was dependent on RomR. Specifically, in RomR-depleted cells escaping the bdelloplast, TadZA-msfGFP still formed mobile foci or clouds but these structures frequently occupied non-polar or sub-polar positions and displayed altered dynamics compared to the control (**Figure 5G**). Thus, our results indicate that the Tad ATPase undergoes highly dynamic RomR-dependent spatial redistribution biased toward the invasive pole during the attack phase.

Collectively, our data support a model in which the Tad system is essential to engage *B. bacteriovorus* into prolonged prey attachment and invasion upon initial contact, and whose unipolar assembly is promoted by RomR, possibly via interaction with the TadZ region of the atypical Tad ATPase (**Figure 6**).

**Figure 6:**
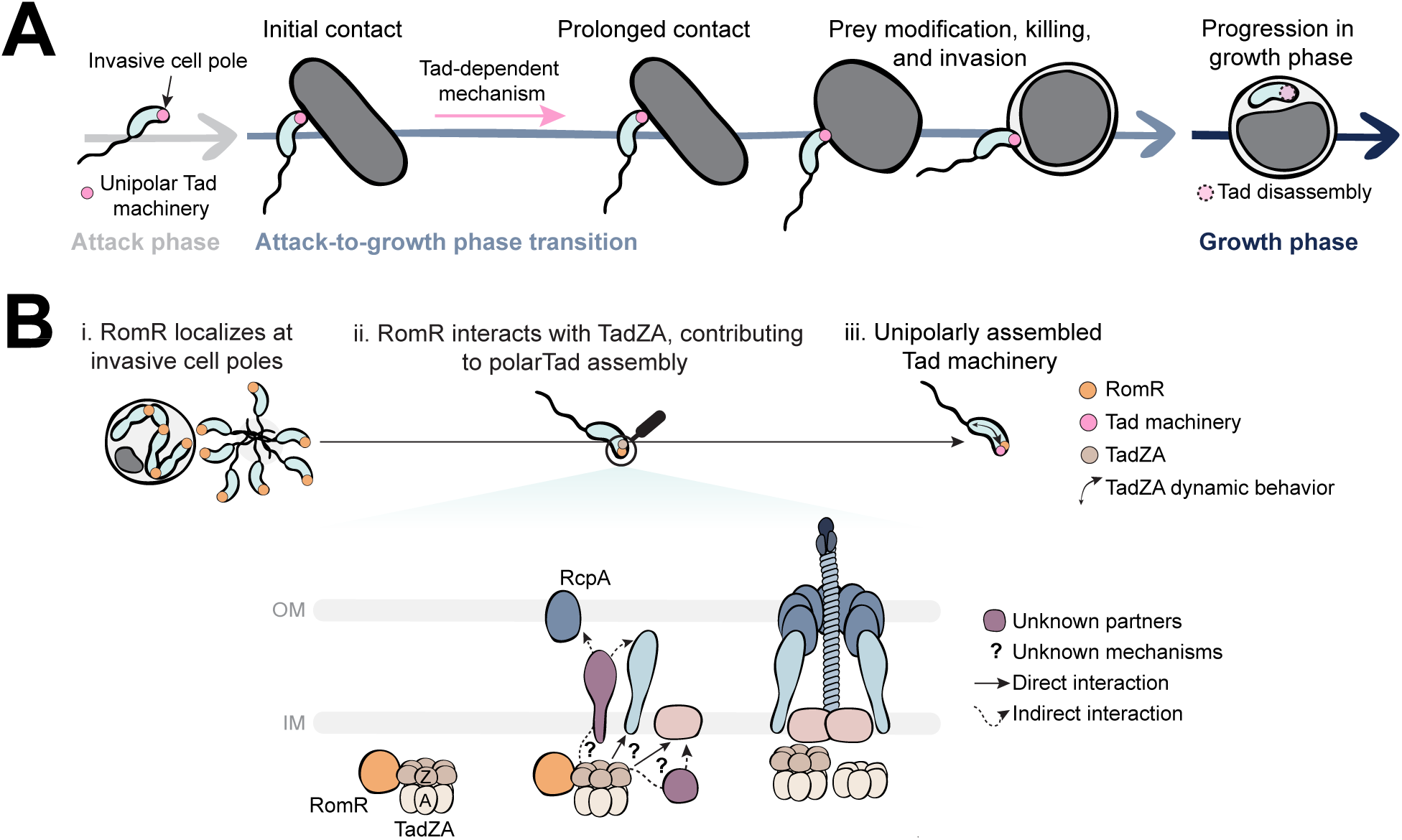
Proposed model for the role, dynamics, and assembly positioning of the Tad machinery in B. bacteriovorus. **A.** The Tad machinery localizes at the invasive cell pole where it is essential for prey attack. In the absence of a functional Tad, prolonged attachment to the prey surface, prey remodeling and prey invasion do not occur, preventing predator cell cycle progression and proliferation. **B.** The essential polar hub protein RomR localizes at the new invasive cell pole where it recruits TadZA through interaction with the TadZ domain and linker region, likely promoting the assembly of the whole Tad machinery at this pole only.

## DISCUSSION

In this study, we identify the Tad machinery as an essential predatory nanomachine coordinated in space and time with the lifecycle of *B. bacteriovorus.* Using conditional CRISPRi combined with live-cell imaging, we show that a complete Tad system is necessary to establish prolonged prey attachment, remodel the prey and proceed to invasion. We further demonstrate that the machinery localizes specifically at the invasive pole of attack-phase cells and that its positioning depends on the polarity factor RomR. Together, these findings reveal the adaptation of a conserved type IV pilus system to the spatial and temporal constraints of obligate bacterial predation.

Our results refine previous indications that the Tad system contributes to the predatory *B. bacteriovorus* lifecycle by clarifying its genomic context, showing essentiality of core components, and providing direct functional analysis in wild-type cells. Imaging in microfluidics enabled for the first time the live visualization of the entire predatory lifecycle at the single-cell level, including the highly dynamic processes of prey attack and invasion that cannot be captured on a microscopy pad. These experiments revealed that Tad-depleted predators still establish transient contacts with prey, indicating that the machinery is not strictly required for initial encounter. However, the observed defects in stable attachment and invasion position the Tad system as a key factor in the transition between reversible prey interaction and productive predation. Whether the Tad machinery or pilus contributes directly to prolonged prey attachment and/or invasion itself remains to be determined. Nevertheless, the absence of detectable alteration to prey cell shape or viability upon transient contact with Tad-depleted *B. bacteriovorus* strongly suggests that Tad activity acts upstream of the cellular program leading to bdelloplast formation. The inability of RomR-depleted cells to establish even transient contact with prey further suggests that other machineries, which may be impaired in the absence of RomR, may play this role. A palette of adhesins was proposed to mediate specific prey recognition ^44^. However, their function remains unexplored *in vivo*. Besides the Tad system, *B. bacteriovorus* encodes two additional TFF systems, a T4aP and a type II secretion system ^25^, representing attractive candidates for early prey interaction roles. Future work is needed to reveal the mechanism responsible for initial prey-predator encounters and whether it is coordinated with Tad function.

Across bacteria, Tad nanomachines have been implicated in adhesion, surface sensing, natural transformation, host colonization, virulence and prey killing ^2^. Our findings reveal a distinct adaptation of the Tad system in predation. Unlike the *M. xanthus* Kil apparatus, which assembles exclusively at prey-contact sites, the *B. bacteriovorus* Tad machinery is pre-positioned at the invasive cell pole before prey encounter. Conversely, its cell cycle-dependent organization is reminiscent of the unipolar Tad system of *C. crescentus*. This supports the view that diversification of spatial control mechanisms contributes to functional repurposing of conserved Tad machineries. Several features of the *B. bacteriovorus* Tad repertoire also point to extensive evolutionary remodeling of this machinery. The system is genetically dispersed across multiple loci, lacks a dedicated TadV prepilin peptidase and instead, appears to share the T4aP-associated PilD enzyme. In addition, several predicted signaling proteins are encoded near *tad* genes – notably *tadC*, which is separated from *tadB* outside the main cluster unlike in other characterized Tad systems. Moreover, the RomR proximity network connects Tad with signaling proteins and T4aP elements ^23^, consistent with its integration into broader regulatory pathways. Altogether, these observations suggest that the *B. bacteriovorus* Tad system has undergone substantial specialization to fulfil a role in an obligate predatory context. Identifying the molecular link between the Tad machinery and putative associated signaling pathways represents attractive future investigation paths.

Our data show that the Tad nanomachine is assembled at the invasive pole of attack-phase cells, prior to prey encounter, whereas it appears absent during growth and re-localizes in daughter cells escaping the prey (**Fig. 6A).** This spatial and temporal coordination with the progression of predation reinforces the view that Tad function can be coupled with specific lifestyles. The sudden disappearance of the RcpA focus upon prey invasion suggests that the secretin – and perhaps the rest of the machinery – may be subjected to active disassembly and/or protein degradation via an unknown predation-dependent mechanism.

A major finding of this work is that localization of the Tad machinery depends on RomR. Previous studies proposed that RomR contributes to a polarity module at the invasive pole, which includes several TFF core components (Tad and T4aP ATPases), proteins participating to polarity establishment with the RomR homolog in *M. xanthus*, and signaling pathways, such as two-component systems and effectors of the secondary messenger cyclic-di-guanosine monophosphate (c-di-GMP). Here, depletion of RomR disrupted polar localization of the Tad secretin RcpA, whereas mislocalized RomR foci redirected RcpA positioning. These observations provide direct evidence that Tad assembly follows RomR-defined polarity cues and support the emerging view that the invasive pole constitutes an organized platform, orchestrated by the hub protein RomR, that spatially coordinates polarity with multiple predatory functions.

Our data identify the atypical TadZA fusion protein as a candidate molecular link between RomR-driven polarity control and Tad assembly. TadZ proteins belong to the ParA/MinD-like superfamily of AAA ATPases involved in spatial organization of diverse cellular cargos ^45^ and contribute to Tad positioning in several bacteria ^13–15^. Here, we show that RomR interacts with the TadZ-linker region of the *B. bacteriovorus* TadZA fusion protein. Interestingly, TadZA displayed highly dynamic localization behavior during the attack phase, rapidly redistributing between poles while remaining biased toward the invasive pole. To our knowledge, this represents the first description of dynamic localization for a Tad-associated ATPase and suggests a distinct mode of spatial regulation in *B. bacteriovorus*. Although the functional significance of these dynamics remains unclear, the unusual TadZA architecture raises the possibility that spatial regulation and ATPase activity are functionally coupled within a single polypeptide. The conserved genomic vicinity of *tadZ* and *tadA* across bacterial Tad systems may have facilitated the emergence of this fusion, which could confer a selective advantage in *B. bacteriovorus* ^1,2^. Whether TadZA interaction with RomR directly provides a spatial cue for Tad assembly remains to be determined.

Notably, the RomR-TadZA association reveals striking parallels with polarity modules controlling Tad systems in other bacteria. In *C. crescentus*, the TadZ homolog CpaE is required for polar assembly of the Tad machinery and depends on the polar landmark protein PodJ for proper localization. PodJ additionally contributes to cell cycle-dependent regulation of Tad pili through recruitment of a histidine kinase controlling pilin accumulation ^11,12^, further illustrating the interconnection between polarity factors, signaling, and Tad assembly.

TadZ proteins typically contain an N-terminal atypical receiver domain (ARD) proposed to mediate protein-protein interactions involved in polar localization and multiprotein complex assembly ^14,46–48^. Remarkably, the TadZ region of *B. bacteriovorus* TadZA lacks this ARD domain (**Figure S7A**), whereas RomR contains a degenerated receiver domain ^21^. This suggests that functions canonically associated with TadZ may instead be distributed between two interacting proteins in *B. bacteriovorus* (**Figure S7B**), highlighting the modularity of spatial organization principles underlying Tad positioning.

Altogether, our findings establish the Tad machinery as an essential determinant of early predation stages in *B. bacteriovorus* and illustrate how a conserved type IV filament system can be repurposed through integration into a polarity network to support specialized bacterial behaviors.

## METHODS

### Bacterial strains and growth conditions

All bacterial strains used in this study listed in **Table S2,** which will be available in the peer-reviewed version of the manuscript. *E. coli* strains were routinely grown at 30 or 37°C with constant shaking in LB medium, except for induction of POLAR fusions (see below), which was performed in M9 salts supplemented with 0.2% casaminoacids (M9+CASA). Molecular DNA cloning was performed in *E. coli* Top10, NEB DH5a or CC118*λpir*. *E. coli* S17-1 λpir was used as donor strain for RP4-dependent conjugation of mobilizable plasmids into *B. bacteriovorus.* POLAR assays and the production of prepilins and prepilin peptidase were conducted in *E. coli* TB28/pAH69 and MG1655, respectively. *E. coli* prey suspensions were obtained from overnight cultures of MG1655 derivatives, carrying an antibiotic resistance cassette for the selection of antibiotic-resistant *B. bacteriovorus* when appropriate, as described in ^49^. *B. bacteriovorus* strains were cultured in the presence of *E. coli* prey at final OD_600_ = 1 in DNB liquid medium (Dilute Nutrient Broth, Becton, Dickinson, and Company) supplemented with 2 µM CaCl_2_ and 3 µM MgCl_2_ salts (DNBs), at 30°C with constant shaking, following the protocol detailed in ^49^. To isolate single clones, *B. bacteriovorus* were grown on double-layer agar plates consisting of a bottom agar layer (15 g/L agar) and top agar layer (7 g/L agar) containing preys at final OD_600_ = 1 ^49^. Plaques were typically observed after 3-5 days of incubation at 30°C. All *B. bacteriovorus* strains were obtained from the wild-type HD100 reference strain (taxon: 26442) ^25^. When appropriate, antibiotics were added in liquid or solid media at the following concentrations: Ampicillin 50 µg/ml (Amp50) or 200 μg/ml (Amp200), Chloramphenicol 15 μg/ml (Cm), Kanamycin 50 μg/ml (Kan), and Tetracycline 7.5 μg/ml (Tet). For gene expression induction, arabinose or IPTG was added to cell cultures at the indicated time and final concentrations.

### Construction of strains and plasmids

Strains and plasmids construction methods are detailed in **Tables S3-4** and oligonucleotide sequences are listed in **Table S5,** which will be available in the peer-review version of the manuscript. To construct plasmids, standard molecular cloning methods were used, and the assembly of PCR-amplified fragments was achieved using the NEBuilder HiFi mix (New England Bio-labs). Constructs were verified by Sanger sequencing and transferred to *B. bacteriovorus* by conjugation as previously reported ^17^. Briefly, the S17-1 donor strain was harvested and washed twice in DNBs before resuspension at a 1:10 ratio of the initial volume and addition of *B. bacteriovorus* recipient at a 0.6:1 v/v ratio. Transconjugants were selected after at least 4 h and maximum overnight incubation by plating cells on selective medium using the double-layer technique. Scarless allelic replacements were performed using a two-step homologous recombination strategy with pK18*mobsacB*-derived suicide vectors ^17,50^ carrying a kanamycine-resistance gene for selection with Kan and the *sacB* gene conferring sucrose sensitivity for plasmid counter-selection with 5% v/v sucrose, yielding either wild-type or modified genotypes. *rcpA, rcpC, tadZA,* and *tadB* deletion attempts invariably yielded wild-type genotype after the second recombination event (screening of at least 72 isolated plaques across 2 independent matings for each construct). Single plaques were isolated for screening, and chromosome modifications or presence of replicative plasmids (pSEVA251 derivatives carrying the low-copy RSF1010 replicon) were confirmed by PCR and DNA sequencing.

### Identification and annotation of tad genes

To identify Tad components in *Bdellovibrio bacteriovorus* HD100, we analyzed the annotated proteome encoded by the RefSeq reference genome, chromosome accession NC_005363.1, assembly accession GCA_000196175.1, last accessed in April 2022. Type IV filament components were detected with MacSyFinder v2.0rc7 ^32^ (default parameters and default thresholds) with the Tad and other TFF model included in TFFscan v1.0rc2 ^1^ model package, with Tad predictions extracted for further analysis. MacSyFinder predictions were used as the initial candidate set for downstream manual annotation and curation. These annotations were complemented with BLASTp ^51^ sequence homology searches, KEGG-based paralogs identification ^52^, InterPro ^53^ analyses of conserved protein domains and architecture, and exploration of putative functional protein association networks using STRING ^54^.

### CRISPRi-mediated gene silencing

Gene silencing was performed in derivatives of the GL2000 CRISPRi starter strain expressing *dcas9* from *Streptococcus pasteurianus* under the control of the IPTG-inducible promoter P*_tac_* (*Bd0063-Bd0064::*P*_dnaK_-lacI*, *Bd1853-Bd1855::*P*_tac_-dcas9*), developed in ^36^. From the 5’- to 3’-end, sgRNAs consist of a variable 20-bp spacer, specific to the target gene, and an invariable dCas9 scaffold. Spacer sequences are complementary to the non-template strain and target either the −10/-35 region between the transcription start site and the start codon, or the coding sequence, and are located just upstream of a protospacer-adjacent motif (PAM) NNGTGA appropriate for the *S. pasteurianus* dCas9. PAM and target sequences were identified for each genomic region of interest using the CHOPCHOP tool ^55^. sgRNAs were cloned into the pCRISP-I vector (pSEVA251-P*_tac_*-*lacI-*P*_BioFab_*) and transferred by conjugation in CRISPRi starter strains. Silencing of selected gene(s) was performed by inducing *dcas9* expression with 200 µM IPTG in overnight or single-growth cycle prey-predator co-cultures, the latter corresponding to a single round of synchronized predation. For single-growth cycle depletion co-cultures, overnight cultures of *B. bacteriovorus* strains were filtered on 1.2 µm syringe filters, mixed with an excess of prey to limit remaining attack phase cells, and incubated for 5h with IPTG before collecting newborn depleted predators by filtration on 0.8 µm syringe filters. Assessment of predation and protein localization was performed through snapshot, time-course, and time-lapse microscopy (see below).

### Killing assays in microplates

Killing assays were carried out as described in ^49^. Overnight *B. bacteriovorus* cultures of interest were filtered through 0.8 μm syringe filters to eliminate residual bdelloplasts. Predator counts, in plaque forming unit (PFU)/ml, were estimated using the SYBR Green-based assay described in ^49^. In each well of a black 96-well plate with transparent bottom, 2.4 × 10^8^ PFU of filtered predator culture were combined with fresh DNBs and *E. coli* prey MG1655 prey at final optical density at 600 nm (OD_600_) = 1. OD_600_ was monitored every 20 min in a Synergy H1 m plate reader (Biotek), with continuous double orbital shaking and temperature set to 30°C. In a single experiment, each strain was tested in technical triplicates in three different wells. Each experiment was performed in biological triplicates. The CuRveR RStudio package was employed to fit curves to a Richard equation, extract *r_max_* and *s* values, and plot data ^49^.

### Processing of prepilins by PilD prepilin peptidase in *E. coli*

Prepilin processing was assessed by producing 6His-tagged prepilins and the PilD prepilin peptidase in *E. coli* MG1655. An optimized version of *pilD,* in which the TTG start codon was replaced by an ATG start codon, was utilized in these assays to improve translation. *prepilA-6his* and *preflp1-6his* were constitutively expressed from pSEVA251 plasmids under the control of P*_nptII_*, while *pilD* and the catalytic mutants *pilD_D120A_* and *pilD_D184A_* were expressed from pBAD18 plasmids, under the control of an arabinose-inducible promoter. pBAD18 and pSEVA251 plasmids were used to co-transform electrocompetent MG1655 cells. To evaluate pilin processing, overnight starter cultures prepared from single colonies were diluted 1:100 in fresh LB medium supplemented with appropriate antibiotics until OD_600_ reached 0.2. From there, *pilD expression* was induced by adding the indicated arabinose concentration (from 0 to 1% w/v) to the culture. After 1h induction, cells were harvested by centrifugation, and proteins were precipitated and concentrated using 10% (v/v) trichloroacetic acid. The resulting pellet was washed twice with acetone and resuspended in 1x SDS-containing sample buffer before Western Blot analysis (see below).

### SDS-PAGE and Western Blot analysis

Protein samples of attack phase *B. bacteriovorus* were prepared by collecting cells from overnight cultures by centrifugation at 13,000 *g* and resuspending the resulting cell pellets in 4x SDS-containing sample buffer at a 1:20 ratio of the initial volume. Growth phase samples were collected from a prey-predator culture – initiated at a 1:1 volume ratio, typically corresponding to a 1:5 predator:prey cell ratio, to minimize free predator cells upon synchronized prey attack ^41^ – at the indicated timepoints and processed as described for attack phase samples. Samples were loaded on NuPage^TM^ Bis-Tris SDS precast polyacrylamide gels (Invitrogen) and ran at 190 V in NuPAGE^TM^ MOPS (Invitrogren, cat n°NP0001) (**Figure S4A-B and S6C-D)** or MES (Invitrogen, cat n°NP0002) (**Figure S1**) SDS running buffer. Protein transfer on nitrocellulose membrane and Western Blotting were performed using standard methods. Ponceau S staining of the whole membrane was conducted to assess total protein loading. JL-8 monoclonal antibody (Takara, cat n°632380, 1/8000 dilution) and α-mCherry polyclonal antibody (ThermoFisher, cat n°PA5-34974 1/1000 dilution) were used as primary antibodies to detect msfGFP and mCherry fusions, respectively. Goat anti-mouse IgG-horseradish peroxidase (HRP) antibody (Sigma, cat n°A4416, 1/5000 dilution) and goat anti-rabbit IgG-HRP antibody (Sigma, cat n°NA934V, 1/10000 dilution) were used as secondary antibodies for JL8 and mCherry, respectively. His-tagged proteins were detected using the 6x-His Tag Monoclonal Antibody HRP conjugate (Invitrogen, cat n°MA1-21315, 1/10000 dilution). Chemiluminescence generated from the reaction of HRP with luminol (Amersham^TM^ ECL^TM^ Prime, Cytiva cat n° GERPN2232) was captured with an Image Quant LAS 500 camera (GE Healthcare). Brightness and contrast of chemiluminescence images were adjusted in Fiji. For growth phase samples, relative protein levels were evaluated by normalizing band intensities with Ponceau S Staining signal of full lanes, measured in Fiji. Results were plotted in Prism 10 (GraphPad) and figures were assembled in Adobe illustrator (Adobe Inc.).

### PopZ-linked polar recruitment (POLAR) assay

Pairwise protein-protein interactions were tested using the POLAR assay, developed in ^43^ and optimized for cytoplasmic protein pairs using a two-step induction and imaging procedure in ^42^. Bait (RomR) and test (TadZA and corresponding truncations) proteins were fused to msfGFP-H3H4 or mScarlet, respectively. The *romR* gene was cloned in pHCL150 plasmid for C-terminal fusion as in ^23^, and expressed under the control of an arabinose-inducible promoter, while the test genes were cloned in pHCL147 or pHCL151 for C-terminal or N-terminal fusions, respectively, and expressed under the control of an IPTG-inducible promoter. Bait and test plasmids were used to co-transform electrocompetent *E. coli* TB28/pAH69. For imaging, overnight starter cultures prepared from single colonies were diluted 1:100 in fresh LB medium until reaching an OD_600_ of 0.2. Then, cells were collected and resuspended in M9+CASA. From this point, expression of prey and test fusions was induced sequentially. Expression of the test fusion was first induced by adding 100 µM IPTG and incubating the culture for 2 h. Then, the bait fusion was induced by supplementing the culture with 0.2% arabinose (w/v) and incubating for an additional 1h. Cells were imaged after each induction step to (i) confirm that the test fusion does not localize at the cell pole in absence (not shown) of bait and (ii) determine if bait and prey proteins interact.

### Live-cell imaging

*B. bacteriovorus* strains were grown overnight with the appropriate *E. coli* MG1655 prey and antibiotics if maintenance of a replicative plasmid was required. Snapshots of attack phase predators were obtained by spotting cells on 1.2% agarose pads prepared in DNBs medium (hereafter referred to as DNBs-agarose pads). For time-lapse or time-course imaging of synchronous predation cycles, *E. coli* MG1655 were grown in LB medium until exponential phase before being harvested and washed twice in DNBs. *B. bacteriovorus* and *E. coli* cells were mixed with a 3:1 to 5:1 volume ratio to limit the presence of non-predated *E. coli* cells. In all synchronous predation imaging experiments, the prey-predator mixing step is considered as time = 0 min. For time-course experiments, the resulting co-culture mix is incubated at 30°C with shaking and samples are spotted on DNBs-agarose pads at desired time-points for imaging. To assess defects in bdelloplast formation upon CRISPRi-mediated gene silencing, the density of predators was estimated by SYBR Green-based labeling as detailed in ^49^ and mixed at 2:1 predator:prey cell ratio with *E. coli* exponential phase cells before imaging at the indicated timepoints. In time-lapse experiments, the prey-predator co-culture was incubated for 1 h to 2 h at 30°C before spotting cells on a DNBs-agarose pad and imaging the same field of view at regular time intervals, as indicated, with the Okolab enclosure temperature set to 27°C. For POLAR assays (see above), *E. coli* TB28 cells were imaged at indicated timepoints by spotting cells on 1.2% agarose pads prepared in M9 salts medium. Early predation phases were monitored by time-lapse imaging in CellASIC ONIX B04A microfluidic bacteria plates (Merck). Exponential phase MG1655 *E. coli* cells, prepared as above, were loaded at 1 psi and immobilized in microfluidic chambers. In our experiments, *E. coli* cells were typically immobilized in the 0.7 µm-height area of the chamber. Then, predators were perfused at 3 psi until the desired density was reached in the chamber. A DNBs flow was maintained at 1 psi throughout the whole imaging process, with the Okolab enclosure temperature set to 27°C. Cells were imaged at regular time intervals, as indicated. Here, the first time-lapse frame, obtained prior predator arrival in the chamber, is defined as time = 0 min. To monitor predation defects upon CRISPRi, newborn depleted predators and appropriate control strain were perfused in a single (**Figure S5D**) or two distinct imaging chambers (**Figure 2 and S3**) with a continuous flow of DNBs supplemented with 200 µM IPTG.

### Image acquisition

Imaging was performed on Nikon Ti2-E fully motorized inverted epifluorescence microscopes (Nikon) equipped with a CFI Plan Apochromat λDM 100x NA = 1.45 Ph3 oil objective (Nikon), a Prime95b sCMOS 25-mm field-of-view camera (Photometrics), and an Okolab enclosure for temperature control. Epifluorescence microscopy was achieved with a Sola SEII FISH LED illuminator (Lumencor) and filter cubes for mCherry (32 mm, excitation 562/40, dichroic 593, emission 640/75; Nikon) and GFP (32 mm, excitation 466/40, dichroic 495, emission 525/50; Nikon). Highly inclined laminated optical sheet (HiLo) illumination was performed at a 59° incident angle with an iLas2 module (Gataca Systems) using 488 nm 150 mW and 594 nm 100 mW lasers for GFP and mCherry, respectively, and a ZT488/594rpc (Chroma) emission filter to collect light. NIS-Ar (Nikon) or Micro-Manager software were used for epifluorescence or HiLo microscopy, respectively. Pixel size was 0.11 µm or 0.074 µm when using the 1.5 x built-in zoom lens of the microscope. Identical illumination settings were applied when imaging several strains and/or conditions in one experiment and across replicates, and were set to the minimum for time-lapse acquisitions to reduce phototoxicity and preserve cell viability. Microfluidics was conducted with a CellASIC ONIX2 microfluidic system (Millipore). Pressure-driven flow into the cell chambers was controlled by the associated software.

### Image processing

Microscopy images were processed with Fiji ^56^ to crop regions of interest, adjust brightness and contrast, and add scale bars. Brightness and contrast settings were kept identical for all images within a figure panel, unless otherwise indicated. When required, timelapse images were aligned on the first phase contrast frame using the Image Stabilizer plugin in Fiji. For **Figures 3B-2C-S3A-S3B,** the Denoise.ai function (NIS, Nikon) was employed to improve the display of time-lapse images acquired with low exposure, required to preserve cell viability. Similarly, in **Figures 3C**, the Smooth Fiji function was applied to phase contrast images. Cell outlines were manually drawn in Adobe Illustrator from phase contrast and fluorescence images. Figure panels were assembled and annotated in Adobe Illustrator.

### Image analysis

Quantitative image analysis was performed on raw fluorescence images with the Fiji MicrobeJ plugin (version 5.13p) ^57^. Cell outlines were automatically detected with the Bacteria function and verified with the manual editing interface. In **Figure 5F**, predators attached to their prey were manually segmented using the manual-editing interface. Diffraction-limited fluorescent foci were automatically detected, localized with subpixel precision, and associated with their parent cell with the Maxima function using the point option. For **Figure 4C, S4F**, detected Maxima were further verified with the manual editing interface. Cell morphology parameters (e.g., cell length or width), mean fluorescence intensities, and maxima counts per cell were retrieved from MicrobeJ results tables and plotted in Prism 10 (GraphPad) or RStudio. 2D heatmaps of maxima subcellular localization along the relative cell length (**Figure 3A**) were obtained in MicrobeJ and oriented, with polarity option, based on the cell pole with the highest mean fluorescence for the selected fluorescence signal. Demographs represent the distribution of fluorescence signal projected along the medial axis of the cell, normalized to its maximum intensity value, with cell length rescaled from 0 to 1 (relative cell length). Kymographs correspond to the distribution of fluorescence signal projected along the medial axis of the cell over time, without normalization, and with cell length rescaled from 0 to 1 (relative cell length). Demographs and kymographs were plotted in MicrobeJ and oriented with polarity option, based on the cell pole with the highest mean fluorescence for the selected fluorescence signal. In **Figure 5D-E-F**, mean fluorescence values of polar segments, defined as the first and last 20% of the relative cell length, were extracted for each time-point to calculate individual and mean log_2_ of pole/pole fluorescence ratios in RStudio, and data were plotted in Prism 10 (GraphPad). Filtering of attack phase cells based on fluorescent signal (**Figure S5C and E**) was achieved with the Type function. Pole-to-pole fluorescence profiles (**Figure 5B**), corresponding to the mean fluorescence intensity along the relative cell length, were retrieved from MicrobeJ and relative mean fluorescence were plotted in Prism 10 (GraphPad) after calculation in Microsoft Excel, by dividing the mean fluorescence value of each segment by the total fluorescence. All plots were included in figure panels and annotated in Adobe Illustrator. In **Figures 2B and S5B**, the number of non-predated *E. coli* cells and bdelloplasts was manually determined using cell roundness as a proxy for bdelloplast formation, at each timepoint and in each condition with the Fiji Cell Counter plugin. Results were plotted in Prism 10 (GraphPad) and a one phase decay fit curve was applied. The recording of specific events from microfluidics experiments, i.e. prey-predator contacts (initiation, end, and duration **Figure 2D-E and Figure S5D**) and RcpA-mCherry focus loss relative to prey contact and entry (**Figure 3B**), was performed manually from phase contrast and epifluorescence images. Results were plotted in Prism 10 (GraphPad) or RStudio. Time-lapse frames of bdelloplast escape and fluorescence focus appearance of **Figure 3D** were manually determined from phase contrast and fluorescence images of microfluidics experiments and plotted in Prism 10 (GraphPad).

## ACKNOWLEDGEMENTS

We are grateful to members of the Laloux lab and Jovana Kaljević for critical reviewing of the manuscript, Charles de Pierpont for constructing the plasmid used in strain GL2872, Agathe Couturier for expert advice with MicrobeJ, and Julien Herrou and Tâm Mignot for insightful discussions about Tad systems.

