## Supplementary Figures and Legends for "A polarity-controlled Tad nanomachine enables prey invasion in a bacterial predator"

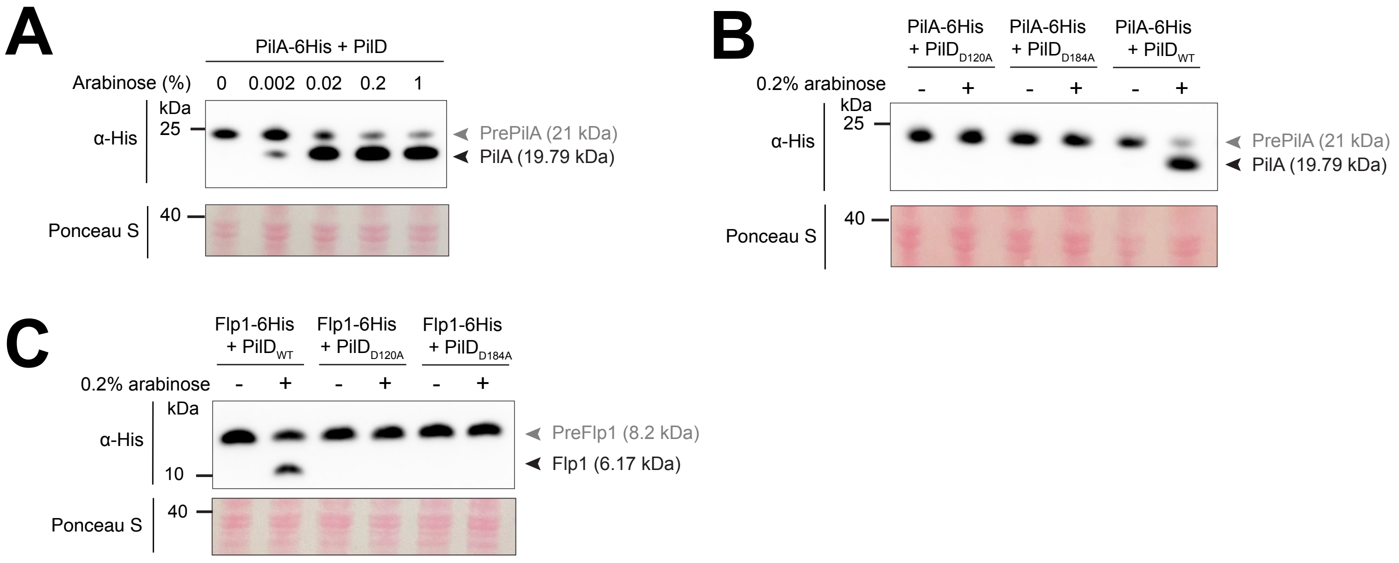
A
PilA-6His + PilD
Arabinose (%) 0 0.002 0.02 0.2 1
kDa
25
α-His
PrePilA (21 kDa)
PilA (19.79 kDa)
40
Ponceau S
B
PilA-6His + PilD$_{D120A}$
PilA-6His + PilD$_{D184A}$
PilA-6His + PilD$_{WT}$
0.2% arabinose - + - + - +
kDa
25
α-His
PrePilA (21 kDa)
PilA (19.79 kDa)
40
Ponceau S
C
Flp1-6His + PilD$_{WT}$
Flp1-6His + PilD$_{D120A}$
Flp1-6His + PilD$_{D184A}$
0.2% arabinose - + - + - +
kDa
α-His
10
PreFlp1 (8.2 kDa)
Flp1 (6.17 kDa)
40
Ponceau S

**Figure S1**

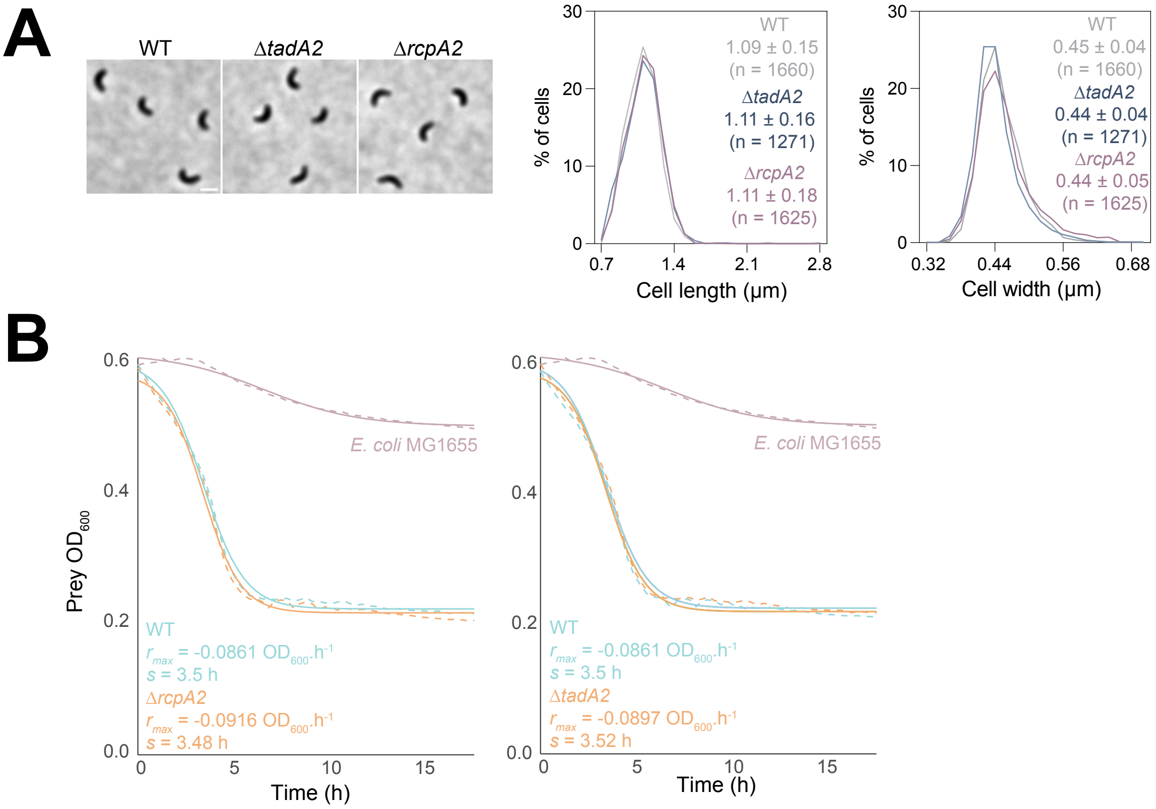
A
WT
ΔtadA2
ΔrcpA2
% of cells
WT
1.09 ± 0.15
(n = 1660)
ΔtadA2
1.11 ± 0.16
(n = 1271)
ΔrcpA2
1.11 ± 0.18
(n = 1625)
Cell length (µm)
WT
0.45 ± 0.04
(n = 1660)
ΔtadA2
0.44 ± 0.04
(n = 1271)
ΔrcpA2
0.44 ± 0.05
(n = 1625)
Cell width (µm)
B
Prey OD600
E. coli MG1655
WT
rmax = -0.0861 OD600.h-1
s = 3.5 h
ΔrcpA2
rmax = -0.0916 OD600.h-1
s = 3.48 h
Time (h)
E. coli MG1655
WT
rmax = -0.0861 OD600.h-1
s = 3.5 h
ΔtadA2
rmax = -0.0897 OD600.h-1
s = 3.52 h
Time (h)

### Figure S2

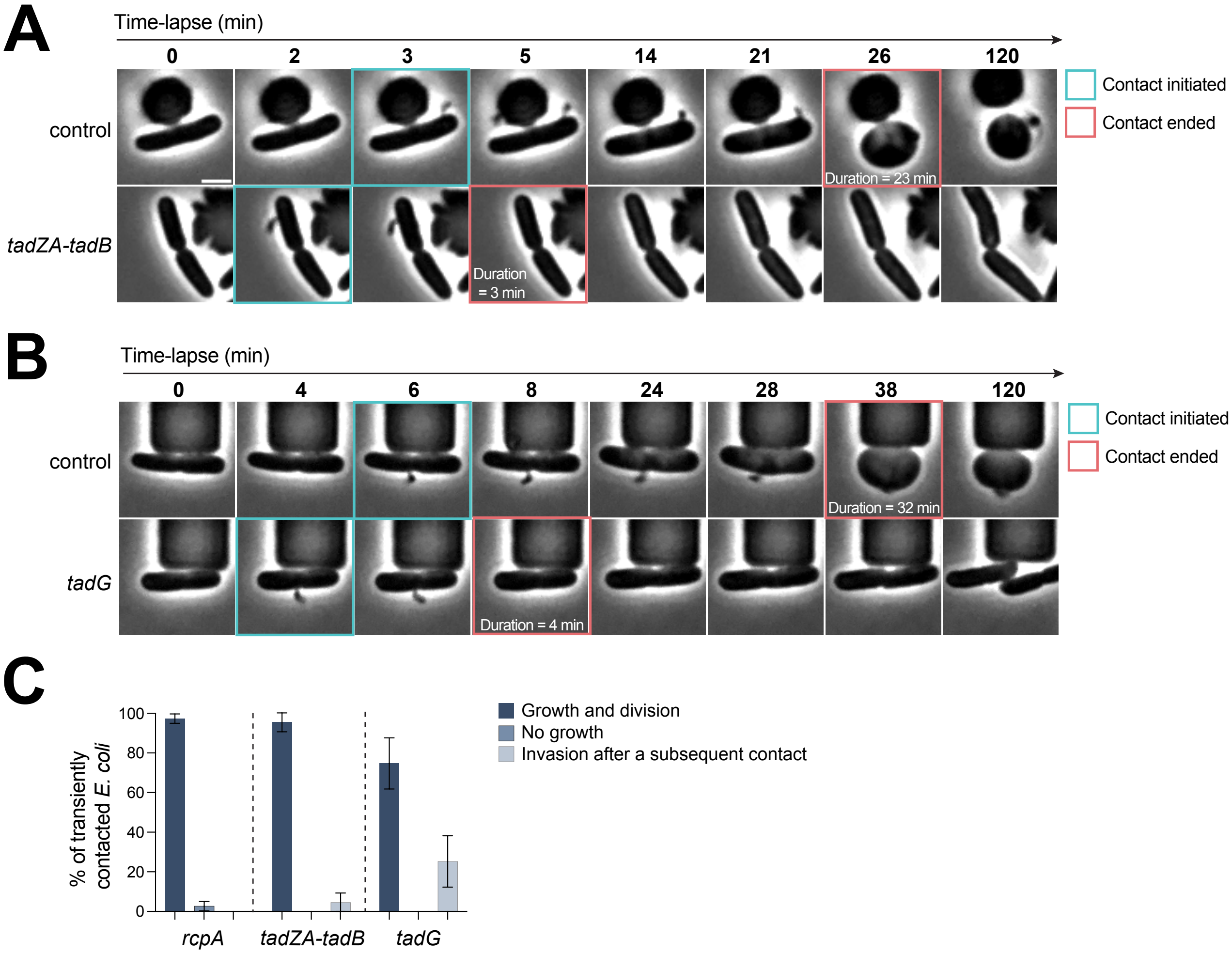
A
Time-lapse (min)
0
2
3
5
14
21
26
120
control
tadZA-tadB
Duration = 23 min
Duration = 3 min
Contact initiated
Contact ended
B
Time-lapse (min)
0
4
6
8
24
28
38
120
control
tadG
Duration = 32 min
Duration = 4 min
Contact initiated
Contact ended
C
% of transiently contacted E. coli
0
20
40
60
80
100
rcpA
tadZA-tadB
tadG
Growth and division
No growth
Invasion after a subsequent contact

### Figure S3

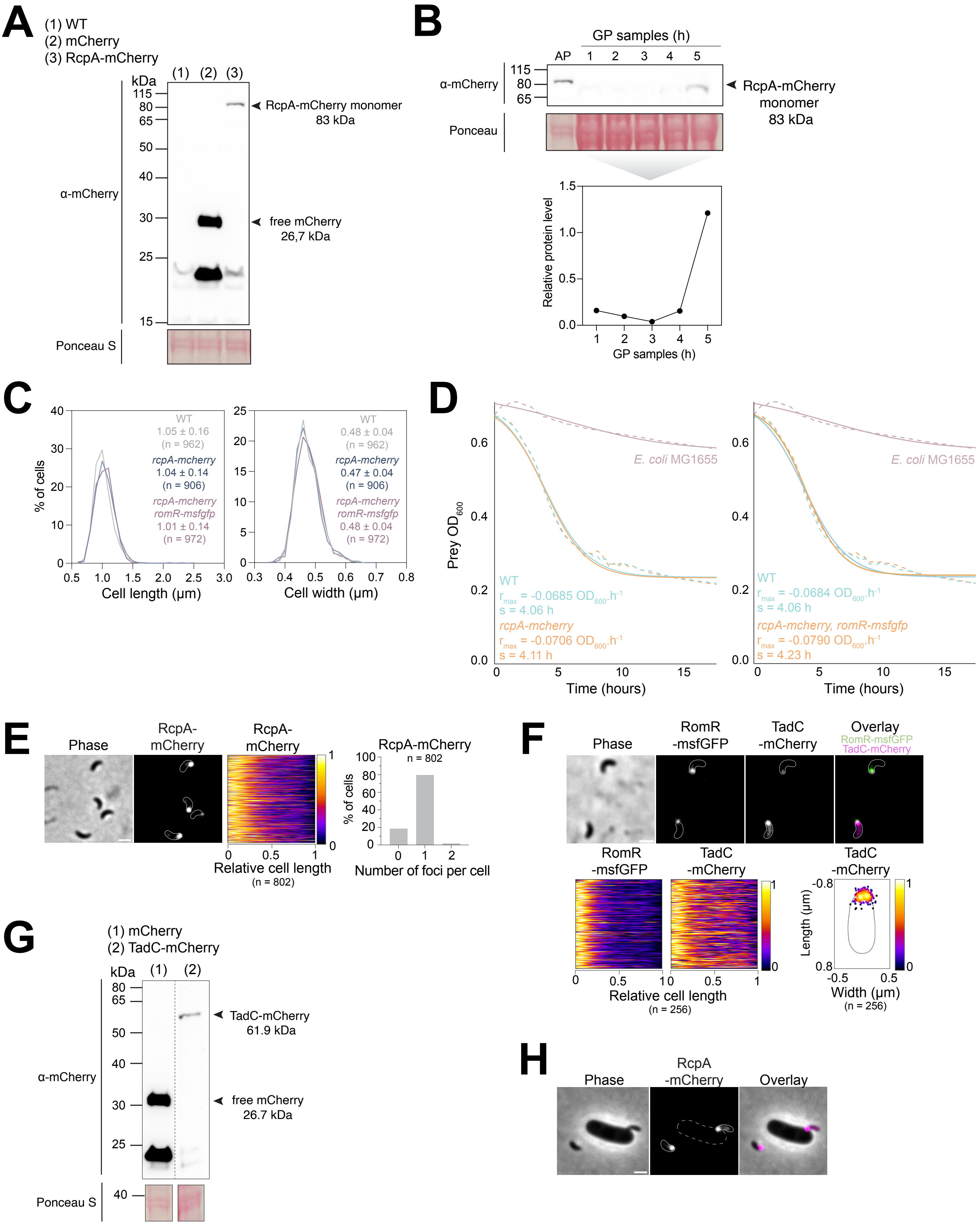
A
(1) WT
(2) mCherry
(3) RcpA-mCherry
kDa
(1) (2) (3)
115
80
65
50
40
30
25
15
α-mCherry
RcpA-mCherry monomer
83 kDa
free mCherry
26,7 kDa
Ponceau S
B
GP samples (h)
AP 1 2 3 4 5
α-mCherry
115
80
65
RcpA-mCherry
monomer
83 kDa
Ponceau
Relative protein level
1.5
1.0
0.5
0.0
1 2 3 4 5
GP samples (h)
C
% of cells
40
30
20
10
0
WT
1.05 ± 0.16
(n = 962)
rcpA-mcherry
1.04 ± 0.14
(n = 906)
rcpA-mcherry
romR-msfgfp
1.01 ± 0.14
(n = 972)
0.5 1.0 1.5 2.0 2.5 3.0
Cell length (µm)
25
20
15
10
5
0
WT
0.48 ± 0.04
(n = 962)
rcpA-mcherry
0.47 ± 0.04
(n = 906)
rcpA-mcherry
romR-msfgfp
0.48 ± 0.04
(n = 972)
0.3 0.4 0.5 0.6 0.7 0.8
Cell width (µm)
D
Prey OD600
0.6
0.4
0.2
0.0
E. coli MG1655
WT
rmax = -0.0685 OD600.h-1
s = 4.06 h
rcpA-mcherry
rmax = -0.0706 OD600.h-1
s = 4.11 h
0 5 10 15
Time (hours)
E. coli MG1655
WT
rmax = -0.0684 OD600.h-1
s = 4.06 h
rcpA-mcherry, romR-msfgfp
rmax = -0.0790 OD600.h-1
s = 4.23 h
0 5 10 15
Time (hours)
E
Phase
RcpA-
mCherry
RcpA-
mCherry
1
0
0 0.5 1
Relative cell length
(n = 802)
RcpA-mCherry
n = 802
% of cells
100
80
60
40
20
0
0 1 2
Number of foci per cell
F
Phase
RomR
-msfGFP
TadC
-mCherry
Overlay
RomR-msfGFP
TadC-mCherry
RomR
-msfGFP
TadC
-mCherry
1
0
0 0.5 1 0 0.5 1
Relative cell length
(n = 256)
TadC
-mCherry
-0.8
0.8
Length (µm)
-0.5 0.5
Width (µm)
(n = 256)
G
(1) mCherry
(2) TadC-mCherry
kDa
(1) (2)
80
65
50
40
30
25
α-mCherry
TadC-mCherry
61.9 kDa
free mCherry
26.7 kDa
Ponceau S
40
H
Phase
RcpA
-mCherry
Overlay

### Figure S4

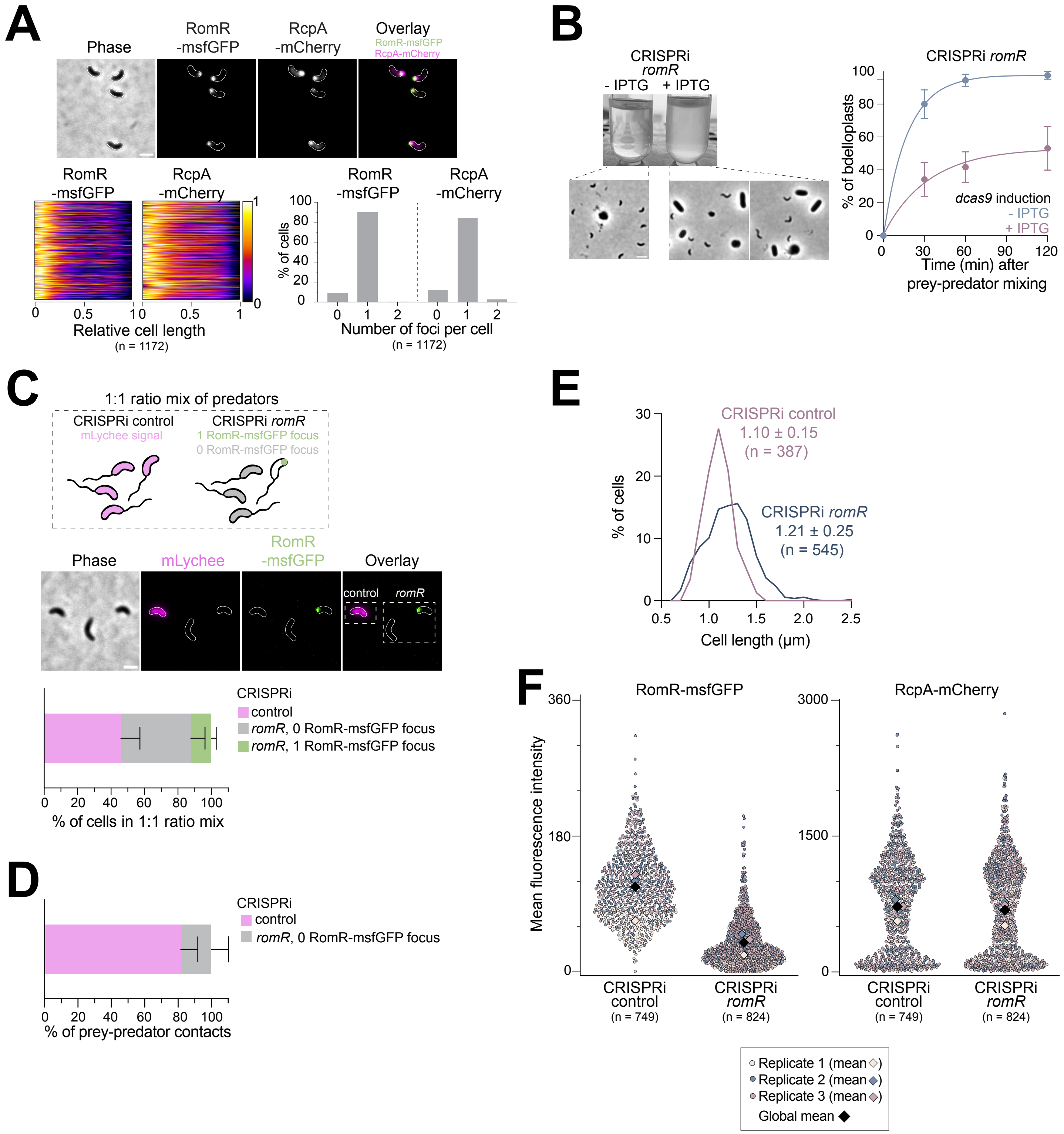
A
Phase
RomR -msfGFP
RcpA -mCherry
Overlay
RomR-msfGFP
RcpA-mCherry
Relative cell length
(n = 1172)
% of cells
Number of foci per cell
(n = 1172)
B
CRISPRi romR
- IPTG + IPTG
CRISPRi romR
% of bdelloplasts
dcas9 induction
- IPTG
+ IPTG
Time (min) after prey-predator mixing
C
1:1 ratio mix of predators
CRISPRi control
mLychee signal
CRISPRi romR
1 RomR-msfGFP focus
0 RomR-msfGFP focus
Phase
mLychee
RomR -msfGFP
Overlay
control
romR
CRISPRi
control
romR, 0 RomR-msfGFP focus
romR, 1 RomR-msfGFP focus
% of cells in 1:1 ratio mix
D
CRISPRi
control
romR, 0 RomR-msfGFP focus
% of prey-predator contacts
E
CRISPRi control
1.10 ± 0.15
(n = 387)
CRISPRi romR
1.21 ± 0.25
(n = 545)
% of cells
Cell length (µm)
F
RomR-msfGFP
RcpA-mCherry
Mean fluorescence intensity
CRISPRi control (n = 749)
CRISPRi romR (n = 824)
Replicate 1 (mean)
Replicate 2 (mean)
Replicate 3 (mean)
Global mean

### Figure S5

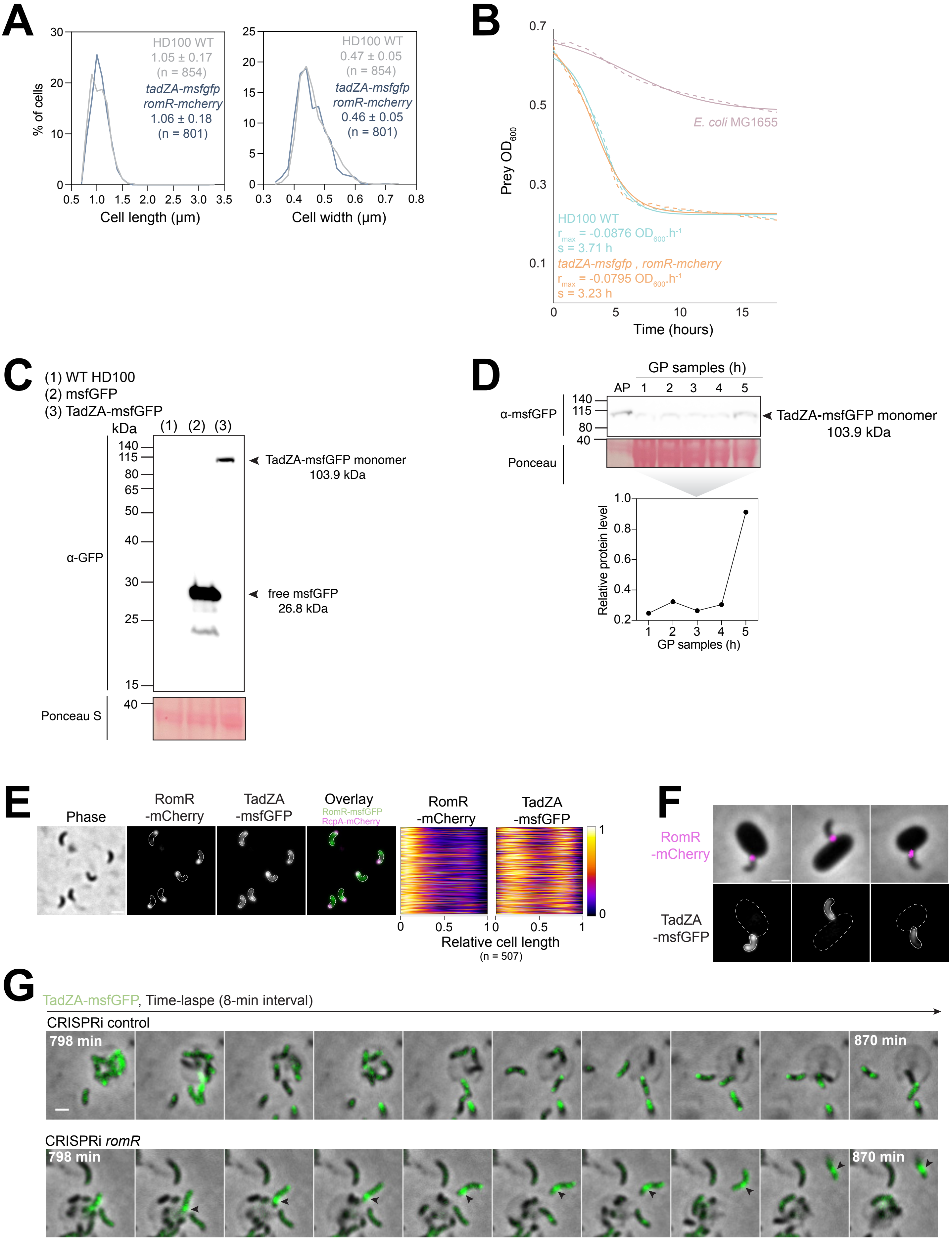
A
% of cells
HD100 WT
1.05 ± 0.17
(n = 854)
tadZA-msfgfp romR-mcherry
1.06 ± 0.18
(n = 801)
Cell length (µm)
HD100 WT
0.47 ± 0.05
(n = 854)
tadZA-msfgfp romR-mcherry
0.46 ± 0.05
(n = 801)
Cell width (µm)
B
Prey OD600
E. coli MG1655
HD100 WT
rmax = -0.0876 OD600.h-1
s = 3.71 h
tadZA-msfgfp , romR-mcherry
rmax = -0.0795 OD600.h-1
s = 3.23 h
Time (hours)
C
(1) WT HD100
(2) msfGFP
(3) TadZA-msfGFP
kDa
α-GFP
TadZA-msfGFP monomer 103.9 kDa
free msfGFP 26.8 kDa
Ponceau S
D
GP samples (h)
AP
α-msfGFP
TadZA-msfGFP monomer 103.9 kDa
Ponceau
Relative protein level
GP samples (h)
E
Phase
RomR -mCherry
TadZA -msfGFP
Overlay
RomR-msfGFP
RcpA-mCherry
Relative cell length
(n = 507)
F
RomR -mCherry
TadZA -msfGFP
G
TadZA-msfGFP, Time-laspe (8-min interval)
CRISPRi control
798 min
870 min
CRISPRi romR
798 min
870 min

### Figure S6

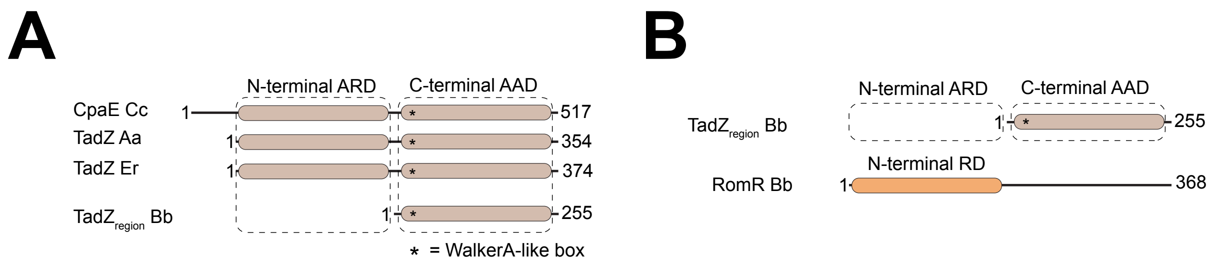
A
N-terminal ARD
C-terminal AAD
CpaE Cc
1
517
TadZ Aa
1
354
TadZ Er
1
374
TadZregion Bb
1
255
* = WalkerA-like box
B
N-terminal ARD
C-terminal AAD
TadZregion Bb
1
255
N-terminal RD
RomR Bb
1
368

### Figure S7

**Figure S1: The T4aP-associated PilD prepilin peptidase processes both T4aP and Tad major pilins.**
Prepilin processing by the PilD prepilin peptidase was assessed in *E. coli* MG1655. **A.** PilD cleaves prePilA, the major pilin of the T4aP. Anti-His ($\alpha$-His) Western Blot analysis of *E. coli* constitutively producing prePilA-6His and PilD wild-type ($PilD_{WT}$; strain GL2194). Cells were grown with increasing amounts of arabinose to induce *pilD* expression. **B.** Mutagenesis of key residues in the PilD active site abolish prePilA maturation. a-His Western Blot analysis of *E. coli* producing prePilA-6His and $PilD_{WT}$, $PilD_{D120A}$ or $PilD_{D184A}$ (strains GL2194, GL2241, GL2242, respectively). $PilD_{D120A}$ and $PilD_{D184A}$ are catalytic mutants lacking the two aspartate residues universally conserved in TFF prepilin peptidases (ref Pelicic 2023). Cells were grown in absence (-) or presence (+) of 0.2% arabinose to induce the expression of *pilD* and corresponding mutants. **C.** $PilD_{WT}$, but not the catalytic mutants, can process preFlp1, a major pilin of the Tad system. a-His Western Blot analysis of *E. coli* cells producing preFlp1-6His and $PilD_{WT}$, $PilD_{D120A}$ or $PilD_{D184A}$ (GL2240, GL3563, GL3564, respectively). Cells were grown in absence (-) or presence (+) of 0.2% arabinose to induce the expression of *pilD* and corresponding mutants. **A-C.** Grey and black arrowheads indicate pre- or processed pilins, respectively. Expected molecular weights (kDa) are indicated on each panel. Ponceau S staining of membranes are shown as loading and transfer control.

**Figure S2: *tadA2* and *rcpA2* are dispensable for predation.**
**A.** *B. bacteriovorus* cells lacking *rcpA2* or *tadA2* display normal cell morphology. Left: representative phase contrast images of attack phase cells of wild-type (WT), *ΔtadA2*, and *ΔrcpA2* strains. Right: histograms of cell length and width of WT, *ΔtadA2*, and *ΔrcpA2* strains (GL734, GL3374, and GL3373, respectively). Mean and standard deviation values, as well as the number of cells analyzed (n) in a representative experiment from two independent biological replicates, are indicated on corresponding graphs. Scale bar = 1 µm. **B.** Genetic inactivation of *rcpA2* or *tadA2* has no impact on predation efficiency. Representative killing curves showing the decrease in optical density at 600 nm ($OD_{600}$) of *E. coli* MG1655 mixed with *ΔrcpA2* (left; GL3373) or *ΔtadA2* (right; GL3374) *B. bacteriovorus* cells, in comparison to WT. $r_{max}$ values correspond to the killing rate and $s$ values to the time at which $r_{max}$ is reached. Dotted and plain lines represent the mean and fit of technical triplicates, respectively, from a representative experiment. Both *ΔrcpA2* and *ΔtadA2* strains were tested in biological triplicates, in comparison to WT ($r_{max}$ and $s$ values are available in **Table S1,** which will be provided in the peer-reviewed version of the manuscript).

**Figure S3: An intact Tad machinery is required for prolonged prey attachment and invasion**
**A-C.** Tad-associated phenotypes were assessed with CRISPRi strains expressing sgRNAs for the targeted repression of *rcpA* (GL2000 *rcpA*sgRNA), *tadZA-tadB* (GL2000 *tadZA-tadB*sgRNA), and *tadG* (GL2914). For clarity, only the targeted gene is indicated in each panel. The control condition corresponds to a strain expressing a non-targeting sgRNA (GL2915). After a single synchronized predation cycle in presence of 200 µM IPTG to induce *dcas9* expression, newborn Tad-depleted and control predators were perfused in separate microfluidics chambers containing immobilized exponentially grown *E. coli*. Early stages of predation were monitored by time-lapse phase contrast microscopy with 1-min (RcpA and TadZA-TadB depletion) or 2-min (TadG depletion) intervals for 2h. **A.** Representative phase contrast images of prey-predator interactions in control (top) or TadZA-TadB-depleted (bottom) conditions, showing selected timepoints of a time-lapse experiment. Time points at which contact initiates (blue squares) and ends (red squares), as well as the resulting contact duration, are indicated for each condition. **B** Representative phase contrast images of prey-predator interactions in control (top) or TadG-depleted (bottom) conditions, showing selected timepoints of a time-lapse experiment. **C.** Histogram showing the fate of *E. coli* cells after being transiently contacted by Tad-depleted predators (i.e., contact observed for ≥2 consecutive time points but followed by detachment during the time-lapse experiment). Bars and error bars represent the mean and standard deviations of three independent biological replicates. n indicates the number of *E. coli* analyzed in three independent biological replicates.

**Figure S4: The Tad fluorescent reporter RcpA-mCherry is produced as a full-length fusion, does not impact predation, and localizes at a single cell pole.**

**A.** The RcpA-mCherry fusion is complete when produced from the native $P_{flp1}$ promoter at the ectopic locus *Bd0063-Bd0064*. Western Blot analysis with an anti-mCherry antibody ($\alpha$-mCherry) of whole-cell protein extracts of the wild-type (WT) *B. bacteriovorus* HD100 strain (lane (1); GL734), a *B. bacteriovorus* strain constitutively producing mCherry (lane (2); GL1025), and a *B. bacteriovorus* strain producing RcpA-mCherry (lane (3), GL2479). All samples were run on the same gel. Ponceau S staining is shown as loading and transfer control. **B.** The RcpA-mCherry fusion is detected in attack phase only. Protein samples of GL2479 were isolated at different timepoints throughout a synchronized predatory cycle: attack phase (AP) and 1h, 2h, 3h, 4h, and 5h after mixing with prey (growth phase (GP) samples). Top: $\alpha$-mCherry Western Blot analysis of the corresponding samples. Bottom: Relative protein levels of GP samples. Ponceau S staining is shown as loading and transfer control and was used to normalize signal intensity of GP samples (see methods). **C-D.** The production of RcpA-mCherry in a WT (GL2479; indicated as *rcpA-mcherry*) or *romR::romR-msfgfp* (GL2585; indicated as *rcpA-mcherry, romR-msfgfp*) background has no impact on cell morphology (**C**) or predation efficiency (**D**). **C.** Histograms of cell length and width in comparison to the WT strain (GL734). Mean and standard deviation values, as well as the number of cells analyzed (n) in a representative experiment are indicated on the corresponding graphs. **D.** Representative killing curves showing the decrease in optical density at 600 nm ($OD_{600}$) of *E. coli* MG1655 mixed with strains producing RcpA-mCherry (WT background, GL2479 and *romR::romR-msfgfp* background, GL2585) in comparison to WT (GL734). $r_{max}$ values correspond to the killing rate and *s* values to the timepoint at which $r_{max}$ is reached. Dotted and plain lines represent the mean and fit of technical triplicates, respectively. Both strains were tested in biological triplicates, in parallel to a WT strain ($r_{max}$ and *s* values are available in **Table S1,** which will be provided in the peer-reviewed version of the manuscript). **E.** Left to right: representative phase contrast and epifluorescence images of strain HD100 *Bd0063-Bd0064::*$P_{flp1}$*-rcpA-mcherry* (GL2479) in attack phase; demograph showing signal distribution along the relative cell length with orientation based on signal intensity; histogram of the number of RcpA-mCherry foci per cell. n indicates the number of cells analyzed in a representative experiment from three independent biological replicates. **F-G.** The IM platform protein TadC localizes at the invasive cell pole in attack phase. **F.** Top: representative phase contrast and epifluorescence images of a *romR::romR-msfgfp B. bacteriovorus* strain producing TadC-mCherry from a low-copy replicative plasmid under the control of the native $P_{tadC}$ promoter, in which both signals are detected (*romR::romR-msfgfp /* pSEVA251-$P_{tadC}$*-tadC-mcherry*, strain GL2939). Bottom left: demographs showing signal distribution of RomR-msfGFP and TadC-mCherry along the relative cell length in cells oriented based on the RomR-msfGFP signal intensity. Bottom right: 2D heatmap of the subcellular localization of TadC-mCherry foci in cells normalized by the cell length and oriented based on RomR-msfGFP signal intensity, with the strongest RomR-msfGFP signal at the top. n corresponds to the number of cells displaying a TadC-mCherry fluorescent focus analyzed in a representative experiment from three independent biological replicates. **G.** TadC-mCherry is produced as a full-length fusion protein. Western Blot analysis with an anti-mCherry antibody ($\alpha$-mCherry) of whole-cell protein extracts of *B. bacteriovorus* cells constitutively producing mCherry (lane (1); strain GL1025), and a *B. bacteriovorus* strain producing TadC-mCherry (lane (2); GL2939). All samples were run on the same gel. For clarity, sample positions were rearranged, as indicated by the dashed line. Ponceau S staining is shown as loading and transfer control. **H.** RcpA-mCherry localizes at the prey-predator interface. Representative phase contrast and epifluorescence images of strain *Bd0063-Bd0064::*$P_{flp1}$*-rcpA-mcherry* (GL2479) after a 15-min incubation with *E. coli*. **E-F, H.** Scale bars = 1 µm.

**Figure S5: RomR depletion impacts predation and RcpA-mCherry localization**

**A.** RcpA and RomR localize as in wild-type cells in a strain carrying a chromosomal *dcas9* expression construct (*pepN-folE::*$P_{flp1}$*-rcpA-mcherry, romR::romR-msfgfp*; GL3343). Left to right: representative phase contrast and epifluorescence images of GL3343 attack phase cells in absence of dCas9 induction (no IPTG); demographs showing signals distribution along the relative cell length in cells orientated based on RomR-msfGFP signal intensity; histogram of the number of RcpA-mCherry and RomR-msfGFP foci per cell. n indicates the number of cells analyzed in a representative experiment from two independent biological replicates. Scale bar = 1 µm. **B.** Depletion of RomR by CRISPRi (GL2913) results in strong predation defects. Left: pictures of overnight prey-predator co-cultures incubated in absence or presence of 200 µM IPTG for induction of *dcas9* expression, and representative phase contrast images of the corresponding overnight cultures. Scale bar = 2 µm. Right: Quantification of bdelloplast formation over time after mixing newborn *B. bacteriovorus* – obtained after a single round of synchronized growth in absence or presence of 200 µM IPTG for *dcas9* induction – with exponentially grown *E. coli* cells. Phase contrast microscopy snapshots were acquired in a time-course experiment at 0, 30, 60 and 120 min post prey-predator mixing, and *E. coli* roundness was used as a proxy for bdelloplast formation. Dots and errors bars represent the mean and standard deviations of at least three independent biological replicates. Data and number of cells analyzed are available in **Table S1,** which will be provided in the peer-reviewed version of the manuscript. **C-E.** RomR-depleted cells fail to contact prey and display more variable cell lengths. RomR depletion was performed in the *romR::romR-msfgfp* CRISPRi strain carrying a *romR*sgRNA (CRISPRi *romR*; GL3199 *romR*sgRNA) and was mixed at a 1:1 ratio with a control CRISPRi strain constitutively producing mLychee (*pepN-folE::*$P_{BioFab}$*-mlychee*) and expressing a non-targeting sgRNA (CRISPRi control; GL3569). For both strains, newborn depleted predators were obtained upon a single synchronized predation cycle with 200 µM IPTG to induce *dcas9* expression. **C.** Microscopy imaging of the generated 1:1 ratio. Top: cartoon representation of the 1:1 ratio mix of newborn predators. Middle: representative phase contrast and epifluorescence images. mLychee-positive cells are the CRISPRi control strain; mLychee-negative cells correspond to the CRISPRi *romR* (with few cells still displaying a RomR-msfGFP focus due to incomplete depletion). Bottom: stacked bar graph of the percentages of CRISPRi control and *romR* cells, further discriminated based on the number of RomR-msfGFP foci they display (0 or 1). The graph presents the mean and standard deviation of three independent biological replicates (n = 932 cells). Scale bar = 1 µm. **D.** RomR-depleted predators rarely contact *E. coli* cells in microfluidics. The 1:1 ratio mix of newborn predators, evaluated in **C**, was perfused in a single microfluidics chamber containing immobilized exponentially grown *E. coli*. Early stages of predation were monitored in time-lapse with 1-min intervals for 2 h. Stacked bar graph showing the proportion of contacts (i.e. observed ≥ 2 consecutive time points) achieved by the CRISPRi control strain or CRISPRi *romR* strain lacking a detectable RomR-msfGFP focus. CRISPRi *romR* cells displaying a RomR-msfGFP focus were discarded from analysis (n = 3 cells in three independent biological replicates) as these correspond to no (or incomplete) RomR depletion. The graph presents the mean and standard deviation of three independent biological replicates (n = 63 contacts). **E.** RomR-depleted cells display more variable cell lengths. Histogram of cell length of CRISPRi *romR* in comparison to CRISPRi control. Mean and standard deviation values, as well as the number of cells analyzed (n=) in three independent biological replicates are indicated. **F.** RomR depletion results in a decrease in RomR-msfGFP, but not RcpA-mCherry, fluorescence intensity, indicating that the loss of RcpA-mCherry focus upon RomR depletion is not due to lower RcpA-mCherry protein amounts. The mean fluorescence intensity of RomR-msfGFP and RcpA-mCherry signals was calculated in the CRISPRi control strain (GL3343 control sgRNA) or the CRISPRi *romR* strain (GL3343 *romR*sgRNA), grown for a single synchronized cycle in presence of 200 µM IPTG to induce *dcas9* expression. Data are shown as scatter plots with colors corresponding to individual biological replicates. The mean per replicate, global mean, as well as the total number of cells analyzed (n =) are indicated.

**Figure S6: The TadZA-msfGFP fusion is dynamic and does not impact predation.**

**A-B.** The production of TadZA-msfGFP (*pepN-folE::*$P_{flp1}$*-tadZA-msfgfp*) in a *romR::romR-mcherry* background (strain indicated as *tadZA-msfgfp, romR-mcherry*; GL3452) has no impact on cell morphology (**A**) or predation efficiency (**B**). **A.** Histograms of cell length and width in comparison to the WT strain (GL734). Mean and standard deviation values, as well as the number of cells analyzed (n=) in a representative experiment are indicated on the corresponding graphs. **B.** Representative killing curves showing the decrease in optical density at 600 nm ($OD_{600}$) of *E. coli* MG1655 preys mixed with strain GL3452 in comparison to WT. $r_{max}$ values correspond to the killing rate and *s* values to the timepoint at which $r_{max}$ is reached. Dotted and plain lines represent the mean and fit of technical triplicates, respectively. This strain was tested in biological triplicates, in parallel to a WT strain ($r_{max}$ and *s* values are available in **Table S1,** which will be provided in the peer-reviewed version of the manuscript). **C.** The TadZA-msfGFP fusion is complete when produced from the $P_{flp1}$ native promoter at the ectopic locus *pepN-folE*. Western Blot analysis with an anti-GFP antibody (α-GFP) of whole-cell protein extracts of the wild-type (WT) *B. bacteriovorus* HD100 strain (lane (1); GL734), a *B. bacteriovorus* strain producing msfGFP from a replicative plasmid (lane (2); GL3454), and a *B. bacteriovorus* strain producing TadZA-msfGFP (lane (3); GL3452). Ponceau S staining is shown as loading and transfer control. **D.** The production of TadZA-msfGFP is cell cycle-regulated. Protein samples of GL3452 were isolated at different timepoints throughout a synchronized predation cycle: attack phase (AP) and 1h, 2h, 3h, 4h, and 5h after mixing with prey (growth phase (GP) samples). Top: α-GFP Western Blot analysis of corresponding samples. Bottom: Relative protein levels of GP samples. Ponceau S staining is shown as loading and transfer control and was used to normalize signal intensity of GP samples (see methods). **E.** TadZA-msfGFP localization pattern is heterogenous in snapshot microscopy images of attack phase cells. Left: representative phase contrast and HiLo fluorescence microscopy images of strain GL3452. Right: demographs showing signals distribution along the relative cell length for cells oriented based on RomR-mCherry signal intensity. **F.** TadZA-msfGFP localization pattern remains heterogenous at prey attachment. Representative phase contrast and HiLo fluorescence microscopy images of GL3452 cells attached to prey after a 20-min incubation with *E. coli*. **G.** TadZA-msfGFP localization is disturbed in newborn RomR-depleted progenies. Overlays of phase contrast and fluorescence images from representative progeny escape obtained as in **Fig. 5G**. Black arrowheads point to disturbed TadZA-msfGFP localization and dynamics (subpolar accumulations rather than polar foci). In panels **E**, **F** and **G**, scale bars = 1 µm

**Figure S7: The domain architecture of the TadZ region in the *B. bacteriovorus* TadZA diverges from described TadZ proteins.**

**A.** The TadZ region of TadZA lacks the N-terminal ARD domain. Schematic representation of the domain architecture of TadZ (CpaE) proteins from *Caulobacter crescentus* (Cc), *Aggregatibacter actinomycetemcomitans* (Aa), and *Eubacterium rectale* (Er), in comparison to $TadZ_{region}$ in *B. bacteriovorus* TadZA (Bb). ARD = atypical receiver domain, AAD = atypical ATPase domain. **B.** RomR displays an N-terminal receiver domain (RD).
